# Simplified model of intrinsically bursting neurons

**DOI:** 10.64898/2026.03.03.709454

**Authors:** Nikhil X Bhattasali, Lerrel Pinto, Grace W Lindsay

## Abstract

Rhythmic neural activity underlies essential biological functions such as locomotion, breathing, and feeding. Computational models are widely used to study how such rhythms emerge from interactions between neuron-level and circuit-level dynamics. Intrinsically bursting neurons are key components of many central pattern generators (CPGs), yet existing models span a tradeoff between biological realism and practical usability. Biophysical models involve many parameters that are difficult to tune, whereas abstract models often integrate poorly into neural circuit simulations. We propose a simplified model of intrinsically bursting neurons derived from a reduced non-spiking biophysical formulation. The model integrates readily into neural circuits while enabling direct and independent control of bursting characteristics, including duration, amplitude, and shape. We show that the model reproduces single-unit biophysical responses to diverse stimuli as well as circuit-level activity patterns from crustacean and mammalian CPGs. This model provides a practical tool for studying rhythm generation in neural circuits.

## Introduction

Computational models are valuable tools for understanding the nervous system, both at the level of individual neurons and neural circuits (***Torres and Varona, 2012***). One particularly fruitful area for such models has been the study of central pattern generators (CPGs), which are neural circuits that generate rhythmic activity to control a variety of rhythmic behaviors, such as locomotion, respiration, and feeding (***Marder and Bucher, 2001; Bucher et al., 2015; Grillner and El Manira, 2020***). CPGs provide an excellent testbed for investigating interactions between neuron-level and circuit-level dynamics, since they often rely not just on structured synaptic connectivity, but also on neurons with intrinsic bursting dynamics, which act as pacemakers by producing periodic bursts even in the absence of synaptic input (***Bucher et al., 2015***).

Different models of intrinsically bursting neurons support different research goals. Detailed conductance-based spiking models (***Hodgkin and Huxley, 1952; Liu et al., 1998***) capture the ionic mechanisms underlying neuronal excitability, enabling investigation into how these mechanisms produce robust dynamics (***Alonso and Marder, 2020***). Reduced spiking models (***Izhikevich, 2003***) simplify conductance-based models through mathematical methods into low-dimensional dynamical systems, allowing burst spiking to be simulated efficiently with few parameters and integrated into circuits (***Tolmachev et al., 2018***). Reduced non-spiking models (***Van Der Pol, 1926; FitzHugh, 1961; Nagumo et al., 1962; Ermentrout, 1994***) produce slow burst envelope waveforms without fast intra-burst spiking, simplifying integration into circuits that use firing rate representations. Phase oscillator models (***Hopf, 1942; Kuramoto, 1975; Buchli et al., 2006***) abstract the core limit cycle of bursting dynamics into polar coordinates, facilitating theoretical analysis and direct frequency control (***Kopell and Ermentrout, 1988; Ijspeert et al., 2007***).

For simulating firing rate activity patterns in complex biological circuits (***Danner et al., 2017***) or synthesizing circuit-based CPGs for neuromechanical bodies (***Ijspeert, 2008; Yu et al., 2014***), it is desirable to use a simplified model that is computationally efficient, readily integrated into neural circuits through synaptic connections, and easily tunable for desired dynamical characteristics—such as burst duration, duty cycle, amplitude, shape, and input dependence. However, existing models tend to occupy opposite ends of an abstraction spectrum. On one end, biophysical models in both spiking and non-spiking variants depend on experimentally derived parameters or nonlinear polynomial coefficients whose effects on dynamical characteristics are highly co-dependent and difficult to tune; even small parameter changes can dramatically alter dynamics (***Izhikevich, 2007***). At the other extreme, abstract models like phase oscillators employ pulse-coupling or phase-coupling connections, which complicates their integration into circuits with diverse neurons and synapses; moreover, producing biologically realistic burst shape, duty cycle, and input dependence requires impractically complicated transformations from a polar coordinate system (***Buchli et al., 2006***).

In this work, we propose a simplified model of intrinsically bursting neurons that addresses these limitations. The model is an analytically tractable, 2-dimensional dynamical system designed to approximate a biophysically derived, reduced non-spiking model. As such, it preserves the biophysical model’s key dynamics and readily integrates into neural circuits, while enabling direct and independent control of bursting characteristics. In evaluation experiments, the simplified model straightforwardly reproduced single-unit responses of the biophysical model under diverse stimuli, and systematic exploration of its parameter space revealed novel response profiles that are difficult to obtain with the biophysical model. Additionally, the simplified model integrated into neural circuits and reproduced canonical activity patterns from crustacean and mammalian CPGs modeled in prior work using spiking and non-spiking biophysical models.

Overall, we demonstrate that the simplified model can be a practical tool for simulating intrinsic bursting within neural circuits. Its interpretable and controllable parameters enable precise tuning to match experimental observations or produce novel dynamics. Consequently, the model is particularly suitable for applications in biological circuit modeling and neuro-robotics.

## Model

We begin with a *biophysical model* of intrinsically bursting neurons from ***Danner et al. (2017)***. This is a rate-coded, reduced non-spiking model, representing an abstraction over reduced spiking models and detailed conductance-based spiking models. Nevertheless, this biophyiscal model captures key dynamics of bursting neurons and has proven useful for simulating circuits, such as the mammalian locomotor circuit in the spinal cord (***Danner et al., 2017; Kim et al., 2022***).

However, the biophysical model also presents challenges. It consists of nonlinear differential equations describing ionic and synaptic currents, with multiple terms including conductances, reversal potentials, activation and inactivation gate dynamics, and time constants. Many of these terms are voltage-dependent and nonlinear, utilizing various sigmoid and bell curves. As such, the relationship between parameter values and dynamics is difficult to predict, and the model is difficult to tune for desired characteristics, which include active and quiet durations at different stimulus strengths, burst amplitude and shape, state switching delays, and stimulus thresholds for tonically active and silent regimes. Tuning the biophysical model to simultaneously satisfy all desired characteristics is cumbersome, even when aided by an automated search algorithm.

We introduce a *simplified model* that preserves the key dynamics of the biophysical model yet presents more interpretable and controllable parameters. For example, each of the characteristics listed above can be directly controlled through independent parameters. The simplified model is constructed by approximating the biophysical model’s dynamics with analytically tractable piecewise-linear functions and hybrid continuous-discrete rules.

The relationship between the models is best understood though their state space geometry, which is a valuable perspective in dynamical systems approaches to neuronal modeling (***Izhikevich, 2007***). Within this framework, differential equations govern the dynamics of a small set of abstract variables (like membrane voltage and adaptation variables) that together capture a neuron’s state. A phase portrait visualizes the state space, with trajectories tracing how state variables evolve jointly through the state space. A nullcline indicates points where a state variable’s time derivative is zero, with equilibrium points found at the intersection of nullclines. Analysis of nullclines and the location, stability, and bifurcations of equilibrium points can provide important insights about the trajectories of a dynamical system.

In the following subsections, we explain how the biophysical model’s state space geometry (***Figure 1A***) produces its activity (***Figure 1B***). Then, we explain how our simplified model reduces complexity in the state space geometry (***Figure 1C***) while producing similar activity (***Figure 1D***).

**Figure 1.**
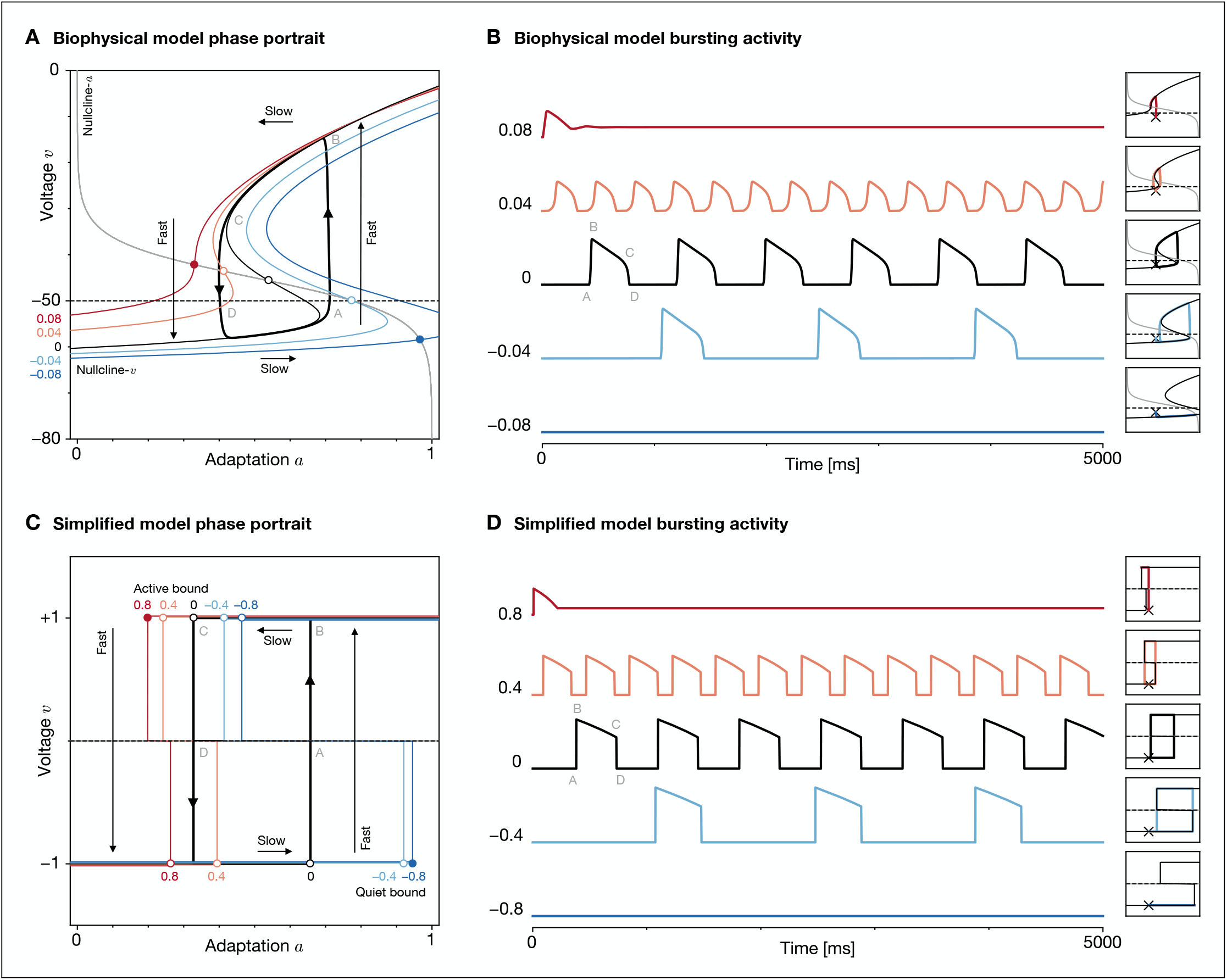
Model dynamics. (A) Phase portrait of the biophysical model, with voltage variable *v* and adaptation variable *a*. Nullclines are shown for various constant stimuli strengths. The firing threshold (dashed line) separates active and quiet states. The system exhibits relaxation oscillations, with slow movement along nullcline-*v* and fast movement across nullcline-*a* to switch between active and quiet states. At strongly table equilibrium (open circle) bifurcates into a stable equilibrium (filled circle) corresponding to tonic and silent modes, respectively. (**B**) Activity of the biophysical model across stimulus strengths, with corresponding phase portrait trajectories. As strength increases, the neuron transitions from the stable silent mode to the periodic bursting mode to the stable tonic mode. (**C**) Phase portrait of the simplified model, with analogous voltage and adaptation variables. Nullclines are piecewise-linear “S”-shaped functions. Trajectories 1) and quiet (*v* = −1) legs until reaching the active and quiet bounds to trigger a fast state switch. The corner ds are unstable in bursting mode and stable in tonic and silent modes. (**D**) Activity of the simplified model responding phase portrait trajectories. Burst shape is controlled through a firing function *y*(*v, a, x*) that input *x* into a firing rate, with a linear-sloped shape used here to approximate the biophysical model. The features of the activity and phase portrait of the biophysical model.

### Biophysical model

The biophysical model is described as a 2-dimensional dynamical system with voltage variable *v* and adaptation variable *a*. The voltage variable (***Equation 1***) represents the neuron’s average membrane potential, which is related to membrane capacitance *C* and various trans-membrane ionic currents *i*. The adaptation variable (***Equation 2***) represents the neuron’s excitability, which the neuron depletes when active and recovers when quiet, and it evolves according to voltage-dependent functions for the steady-state adaptation *a*_∞_(*v*) and time constant τ_*a*_(*v*).

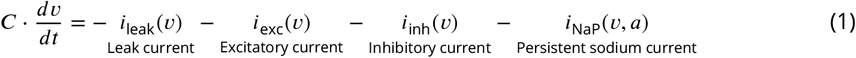

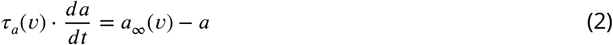

The full description of the model is provided in ***Methods—Biophysical model***.

In the model’s phase portrait (***Figure 1A***), the voltage nullcline is an “S”-shaped function with two knees where trajectories switch between active and quiet states, while the adaptation nullcline is a sigmoid curve that separates regions of depleting and recovering adaptation. Due to differences in the time constants of the state variables, the system exhibits relaxation oscillations during which a trajectory moves slowly along the voltage nullcline’s active or quiet leg, then falls off the knee and switches quickly to the opposite leg across the adaptation nullcline and firing threshold.

At baseline input, the equilibrium point is unstable, and trajectories converge to a limit cycle attractor corresponding to operation in bursting mode. At different input values, the voltage null-cline shifts and stretches in shape, leading to different active and quiet durations (***Figure 1B***). At strongly positive or negative inputs, the unstable equilibrium bifurcates into a stable equilibrium, and trajectories converge to a point attractor corresponding to tonic or silent modes, respectively.

Additional subtle characteristics can be observed by changing input dynamically. First, when changing from silent to bursting mode, there is a delay corresponding to the time for a trajectory starting at the point attractor to move vertically and cross threshold. Such state switching delay is also apparent in bursting mode as the time between falling off a knee and crossing threshold. Second, when changing input while remaining in either stable mode (for instance, changing from strongly negative to weakly negative input while remaining in silent mode), a trajectory may either move to the new stable point or effectively fall off a knee to trigger a state switch. The latter is known as a rebound, which is non-periodic and terminates when the trajectory returns to the new stable point after a single burst. A trajectory’s location during the input perturbation determines which case applies, with a implicit separatrix demarcating the rebound and non-rebound regions.

Finally, the model’s firing activity is a thresholded and scaled function of the voltage variable *v*. As such, the activity waveform’s burst amplitude and shape is closely related to the shape of the voltage nullcline and the rate at which it is traversed (***Figure 1B***).

### Simplified model

The simplified model approximates the biophysical model as a 2-dimensional dynamical system with analogous voltage variable *v* and adaptation variable *a*. In the model’s phase portrait (***Figure 1C***), the voltage nullcline is simplified as a piecewise-linear “S”-shaped function composed of horizontal and vertical segments. A trajectory moves horizontally along the voltage nullcline’s active or quiet leg (*v* = ±1) until it reaches a bound, where it turns to move vertically towards the threshold (*v* = 0); upon reaching it, the trajectory instantaneously switches to the opposite leg.

Such dynamics can be constructed through hybrid continuous-discrete rules. The adaptation variable (***Equation 3***) depletes towards 0 in the active state and recovers towards 1 in the quiet state, with adaptation time constant 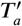, scaling factor *K*_*a*_(*x*), and bounds *a*_active_(*x*) and *a*_quiet_(*x*), which are dependent on net input *x*. The adaptation evolves with an exponential trajectory, which approximates the slow variable trajectory in an averaged burst spiking model (***Izhikevich, 2007***):

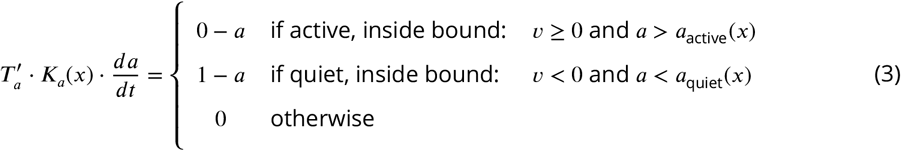

The voltage variable (***Equation 4***) evolves based on a direction function *d*(*v, a, x*) ∈ {−1, 0, +1} and rate functions *r*_active_(*x*) and *r*_quiet_(*x*), with instantaneous switches (of infinite voltage rate) handled by continuously constraining *v* to the range [−1, +1]:

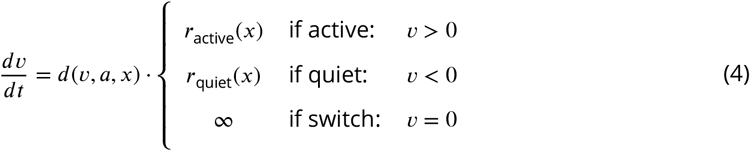

For conciseness, these equations do not include all features, such as stable points, state switching delay, and noise. The full description of the model is provided in ***Methods—Simplified model***.

At different input values, the active and quiet bounds shift, which changes the active and quiet durations. The core mechanism of the model is analytically computing the bounds (***Equation 20***) to achieve desired state durations and input-dependent scaling (***Figure 2A, Equation 24***). This is possible since adaptation *a* is governed by a piecewise-linear and separable differential equation.

**Figure 2.**
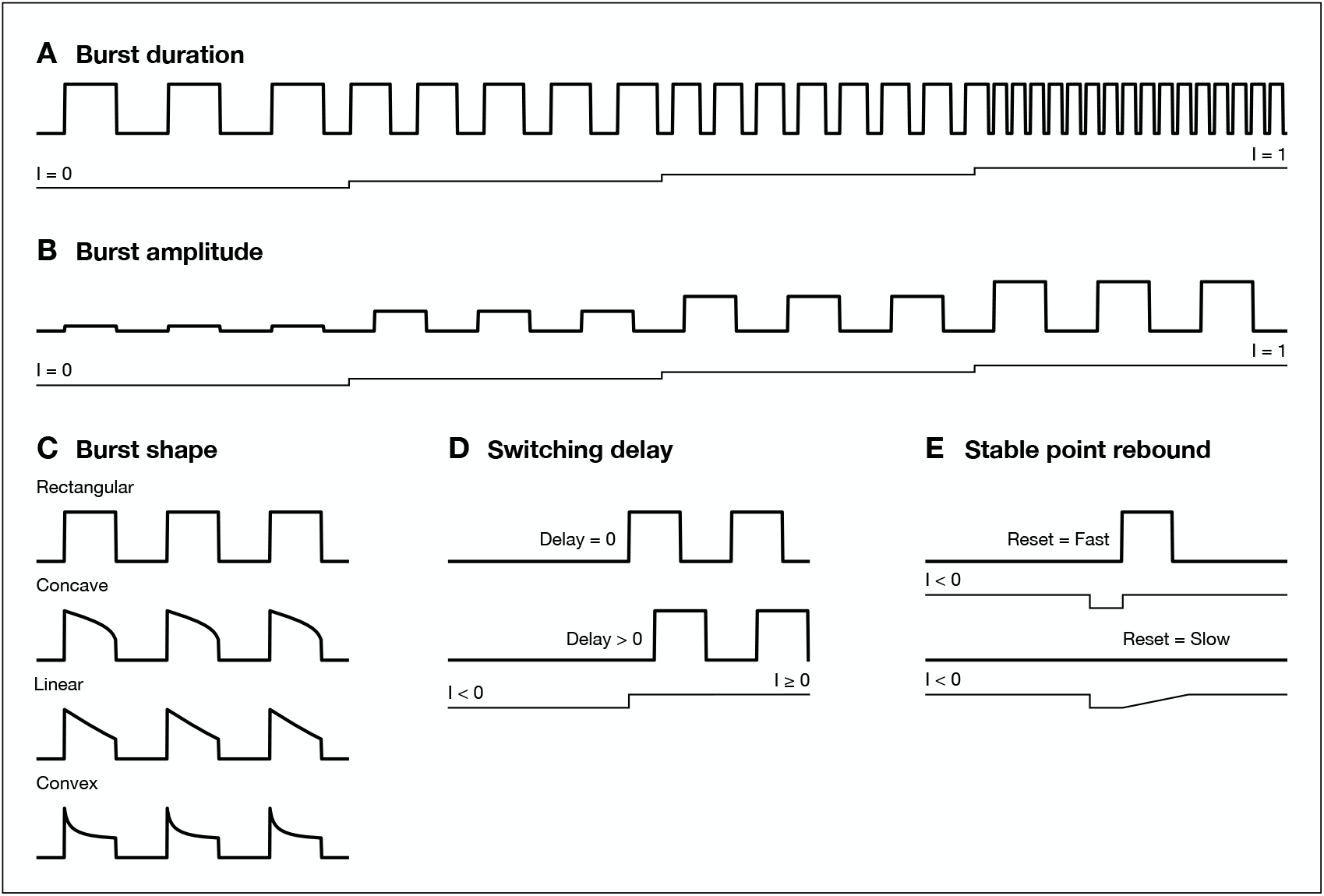
Model features. The simplified model enables explicit control of the bursting waveform through its interpretable parameters. (**A**) Burst duration is controlled through the intrinsic active duration *T*_active_ and quiet duration *T*_quiet_ (achieved at input strength *I* = 0) and the active scale 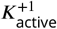 and quiet scale 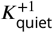 (achieved at *I* = +1). The example shows 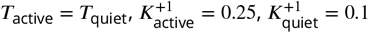. (**B**) Burst amplitude is controlled through the dependence of the firing function *f* on input strength *I*. (**C**) Burst shape is controlled through the shape of the firing function *f* across adaptation *a*. (**D**) Switching delay is controlled by active delay *D*_active_ and quiet delay *D*_quiet_. (**E**) Stable point rebound is controlled by duration scales 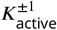 and 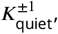 stable point thresholds *X*_active_ and *X*_quiet_, and stable point time margins *δ*_active_ and *δ*_quiet_.

At strongly positive or negative inputs, the corner point of the active or quiet bound becomes stable, transitioning the model to tonic or silent mode (***Figure 1D***). This is achieved by modifying the direction function to reverse trajectories towards the corner point in stable modes (***Equation 29***).

Additional features can be introduced through minor modifications. First, state switching delay (***Figure 2D***) is achieved by modifying the rate functions to increase the time spent on vertical segments of the voltage nullcline (***Equation 27***). Second, stable point rebound (***Figure 2E***) is achieved by introducing a time margin to explicitly control the separatrix between rebound and non-rebound regions around stable points (***Equation 30***).

Finally, the model decouples the shape of active and quiet legs from the activity waveform’s burst amplitude and shape through the firing function *y*(*v, a, x*). The simplest firing function produces a rectangular waveform that oscillates between a high rate *Y*_active_ and low rate *Y*_quiet_:

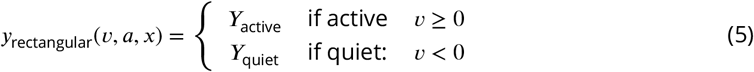

but more complex firing functions enable amplitude modulation (***Figure 2B***) and different burst shapes (***Figure 2C***) while maintaining the same underlying dynamics.

## Results

### Simplified model reproduces single-unit responses of biophysical model

We compared the single-unit responses of the simplified model with those of the biophysical model under 3 types of stimuli—constant, pulse, and periodic—in order to evaluate how well the simplified model captures different properties of the biophysical model. Experiments were run using a consistent set of parameters. For the biophysical model, parameters were used from ***Kim et al. (2022)***, which selected them to match experimental observations of cat locomotion (***Table 1***). For the simplified model, parameters were selected to match the biophysical model (***Table 2***).

**Table 1.** Biophysical model parameters for single-unit experiments.

| Symbol | Name | Value |
| --- | --- | --- |
| $\Delta t$ | Timestep | 1 ms |
| $C$ | Membrane capacitance | 20 pF |
| $G_{\text{leak}}$ | Leak conductance | 4.5 nS |
| $V_{\text{leak}}$ | Leak reversal potential | -62.5 mV |
| $G_{\text{exc}}$ | Excitatory conductance | 10 nS |
| $V_{\text{exc}}$ | Excitatory reversal potential | -10 mV |
| $D_{\text{exc}}$ | Excitatory drive | 0.02 |
| $G_{\text{inh}}$ | Inhibitory conductance | 10 nS |
| $V_{\text{inh}}$ | Inhibitory reversal potential | -75 mV |
| $D_{\text{inh}}$ | Inhibitory drive | 0.00 |
| $G_{\text{NaP}}$ | Sodium conductance | 4.5 nS |
| $V_{\text{NaP}}$ | Sodium reversal potential | 55 mV |
| $V_m$ | Sodium activation midpoint | -40 mV |
| $K_m$ | Sodium activation slope | 6 mV |
| $V_a$ | Sodium inactivation midpoint | -45 mV |
| $K_a$ | Sodium inactivation slope | -4 mV |
| $V_{\tau_a}$ | Sodium inactivation time constant midpoint | -35 mV |
| $K_{\tau_a}$ | Sodium inactivation time constant scale | 15 mV |
| $T_a$ | Sodium inactivation time constant baseline | 320 ms |
| $T'_a$ | Sodium inactivation time constant maximum | 640 ms |
| $V_{\text{threshold}}$ | Firing threshold voltage | -50 mV |
| $V_{\text{max}}$ | Firing maximum voltage | 0 mV |

**Table 2.**
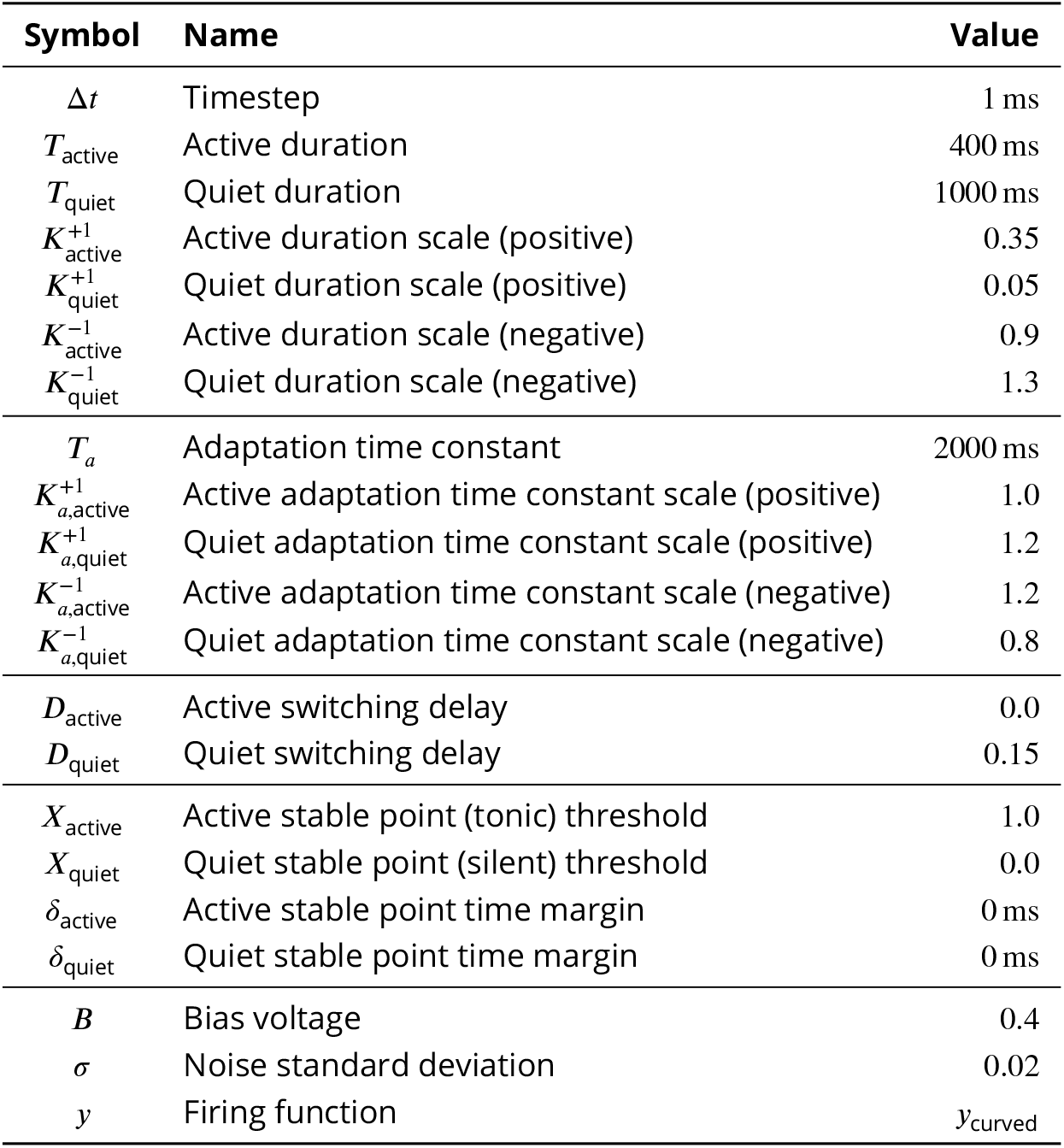
Simplified model parameters for single-unit experiments.

#### Constant stimuli scale the active and quiet durations

To assess duration scaling, we applied constant stimuli of different strengths ranging from strongly positive (excitatory) to strongly negative (inhibitory), and we measured the state durations and duty cycles of waveforms (***Figure 3, Figure 3—figure supplement 1, Figure 3—figure supplement 2***).

**Figure 3.**
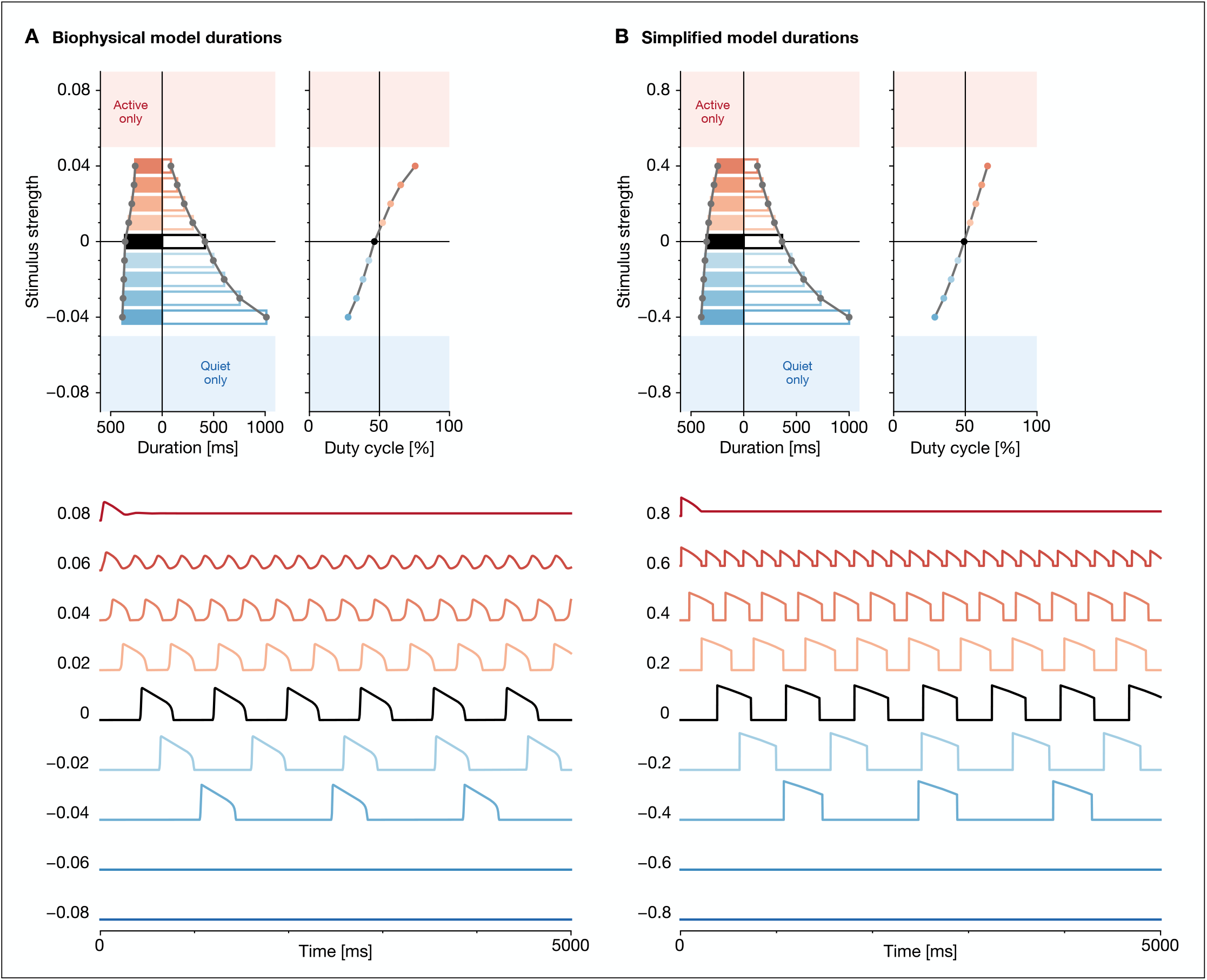
Response to constant stimuli. Top: Durations of the active state (left, filled bars) and quiet state (right, unfilled bars) at various stimulus strengths, along with the resulting duty cycles. Bottom: Activity at representative stimulus strengths. In both the biophysical model (**A**) and simplified model (**B**), the quiet duration decreases faster than the active duration as stimulus strength increases, thus increasing the duty cycle. The models transition into the silent mode (quiet only) at strong negative stimuli and the tonic mode (active only) at strong positive input. **Figure 3—figure supplement 1**. Biophysical model response waveforms to constant stimuli. **Figure 3—figure supplement 2**. Simplified model response waveforms to constant stimuli. **Figure 3—figure supplement 3**. Simplified model effect of parameters on constant response.

In both models, the active and quiet durations decreased with increasing stimulus strengths, with the quiet duration decreasing more rapidly than the active duration. This increased the duty cycle, defined as the fraction of time spent in the active state relative to the total cycle duration. At strongly positive stimuli, both models transitioned into an active-only mode where firing rate oscillated with no quiet state and eventually produced constant tonic firing. At strongly negative stimuli, both models transitioned into a quiet-only mode with no active state. The simplified model closely approximated the duration scaling of the biophysical model. This reflected similarities in the underlying phase portraits of the models, with voltage nullclines shifting and scaling to produce the observed waveform changes (***Figure 3—figure supplement 1, Figure 3—figure supplement 2***). At the same time, there were subtle differences. The simplified model had smooth response profiles, but the biophysical model’s nonlinearities introduced slight non-smoothness for state durations and hence duty cycle. Moreover, the simplified model produced bursts with sharper state switches, which may be reduced by using a firing function with different parameters or a different design.

Additionally, we explored the effect of simplified model parameters on the response to constant stimuli (***Figure 3—figure supplement 3***). The state duration scales could be adjusted to reverse or abolish duration scaling. The adaptation time constant narrowed or widened the duration response profile. The stable point thresholds controlled the input strengths that induced tonic and silent modes. These results suggest parameters to tune for a desired duration response profile.

#### Pulse stimuli shift the oscillation phase

To assess phase shifts, we applied pulse stimuli of different strengths across phases of the intrinsic cycle, which often would advance or delay the waveforms. We quantified phase shift by comparing the perturbed cycle duration to the intrinsic cycle duration, and we plotted finite phase response curves (***Figure 4, Figure 4—figure supplement 1, Figure 4—figure supplement 2***).

**Figure 4.**
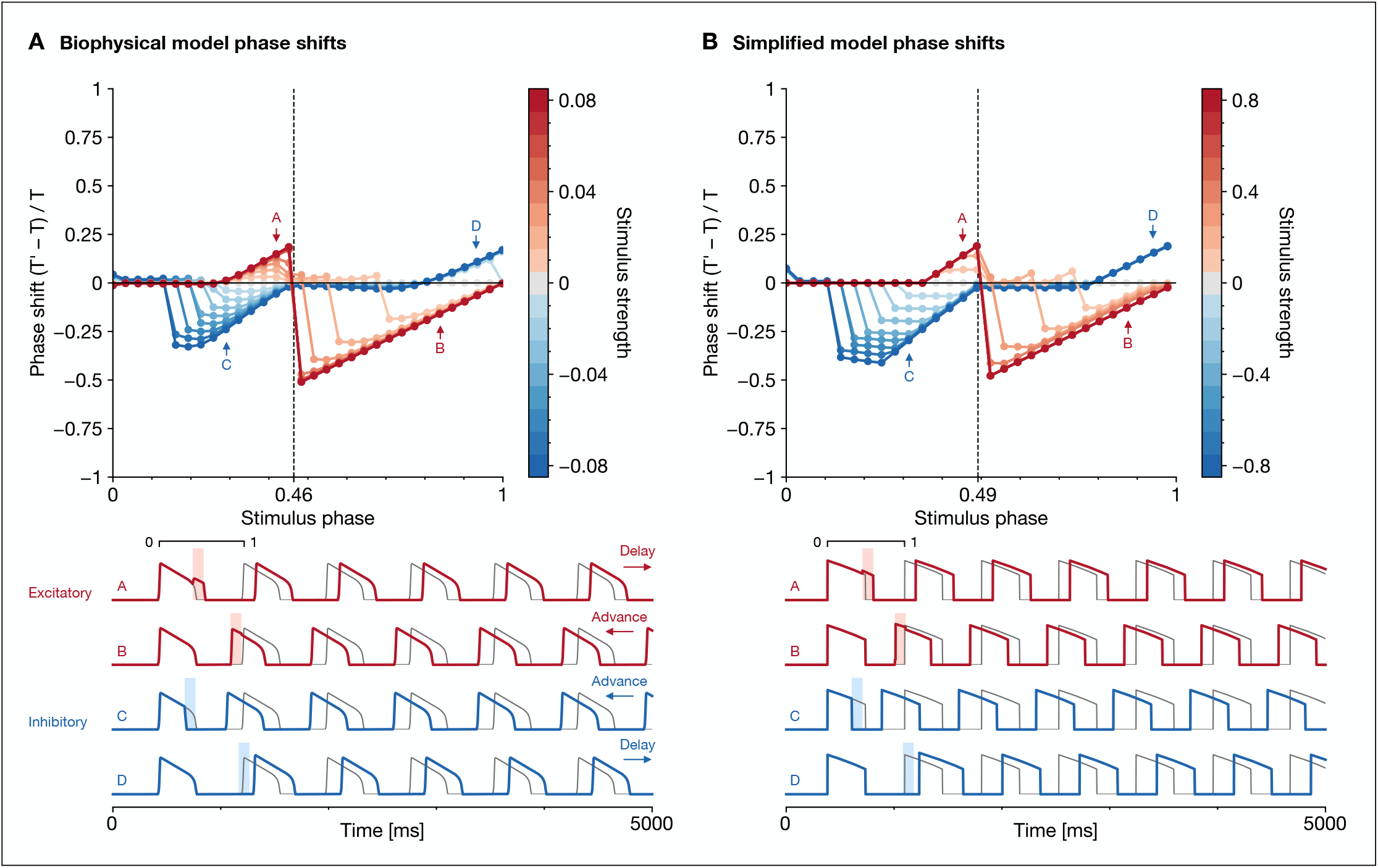
Response to pulse stimuli. Top: Phase response curve showing phase shifts induced by various stimulus strengths and phases (calculated from intrinsic cycle duration *T* and perturbed cycle duration *T* ^′^). The phase of the intrinsic active-to-quiet state switch is marked with a vertical line. Pulse durations are 100 ms. Bottom: Activity of representative responses to excitatory and inhibitory pulses (strengths are the strongest values shown; phases are marked by arrows in the phase response curve). In both the biophysical model (**A**) and simplified model (**B**), pulse stimuli can advance or delay the waveform, depending on the stimulus strength and phase relative to the intrinsic cycle. **Figure 4—figure supplement 1**. Biophysical model response waveforms to pulse stimuli. **Figure 4—figure supplement 2**. Simplified model response waveforms to pulse stimuli. **Figure 4—figure supplement 3**. Simplified model effect of parameters on pulse response.

In both models, excitatory and inhibitory pulses could either advance or delay the waveform depending on their phase relative to the intrinsic cycle. Excitatory pulses tended to lengthen the active state and shorten the quiet state, while inhibitory pulses tended to produce the opposite effects. Stronger pulses of either type produced larger phase shifts. The simplified model closely approximated the phase response curve of the biophysical model, including the locations of flat and diagonal segments and the magnitude of phase shifts. This reflected similarities in the underlying phase portraits of the models. For instance, diagonal segments of the phase response curve corresponded to where trajectories were induced to switch legs of the voltage nullcline, while flat segments corresponded to where trajectories continued on the same leg (***Figure 4—figure supplement 1, Figure 4—figure supplement 2***).

Additionally, we explored the effect of simplified model parameters on the response to pulse stimuli (***Figure 4—figure supplement 3***). The state duration scales determined input-dependent bound shifts, which could dramatically alter the phase response curve. The adaptation time constant affected the distance between active and quiet bounds needed to achieve desired state durations, which in turn altered how easily trajectories switched legs due to pulse perturbations. The adaptation time constant scales resulted in phase-independent speed changes along nullcline legs, which vertically translated flat segments of the phase response curve. The quiet switching delay increased the phase range where firing could be delayed by inhibitory stimuli. These results suggest parameters to tune for a desired phase response curve.

#### Periodic stimuli entrain the oscillation cycle

To assess entrainment, we applied periodic stimuli of different strengths and periods, which often would entrain the waveforms such that firing was phase-locked to the stimulus. We quantified entrainment through the phase coherence of waveform peaks, and we rendered Arnold tongue plots (***Figure 5, Figure 5—figure supplement 1, Figure 5—figure supplement 2***).

**Figure 5.**
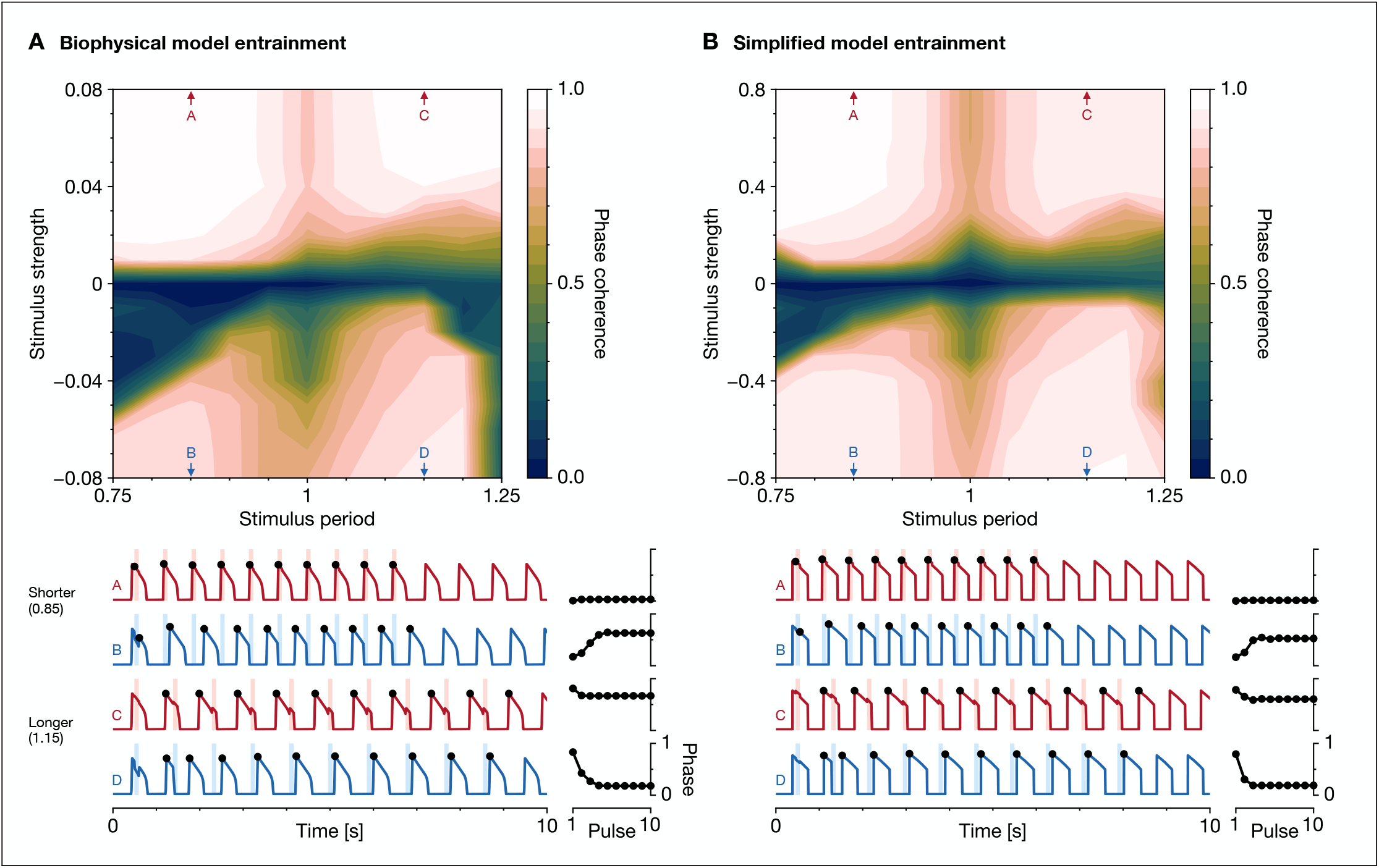
Response to periodic stimuli. Top: Arnold tongue plot showing entrainment level induced by various stimulus strengths and periods (***Arnold and Avez, 1989***). The entrainment level is quantified by phase coherence (calculated from the circular mean of the phases of waveform peaks relative to the stimulus). Pulse durations are 100 ms. Periods are normalized by the intrinsic cycle duration. Bottom-left: Activity of representative responses to pulse trains with shorter and longer periods (strengths and periods are marked by arrows in the Arnold tongue). Bottom-right: Phase of waveform peaks relative to the stimulus, with stabilization of the phase indicating entrainment. In both the biophysical model (**A**) and simplified model (**B**), activity can entrain to positive and negative periodic stimuli. Stronger stimuli more effectively produce entrainment, as do stimuli with periods close to the intrinsic cycle duration. Activity returns to the intrinsic cycle once stimuli are removed. **Figure 5—figure supplement 1**. Biophysical model response waveforms to periodic stimuli. **Figure 5—figure supplement 2**. Simplified model response waveforms to periodic stimuli. **Figure 5—figure supplement 3**. Simplified model effect of parameters on periodic response.

In both models, periodic pulse trains could entrain the waveforms, with more robust entrainment for pulse trains that were stronger or had periods closer to the intrinsic cycle duration. Entrainment was especially poor for shorter-period, inhibitory pulse trains. The simplified model closely approximated the entrainment region of the biophysical model, which reflected similarities in their phase response curves. In particular, the asymptotic locking phases for 1/1 entrainment (***Jiménez et al., 2022***) corresponded to stable points in the phase response curve, which are positive-sloped zero crossings. The models also displayed differences, with slightly stronger entrainment for the biophysical model under excitatory input, and slightly stronger and wider entrainment for the simplified model under inhibitory input.

Additionally, we explored the effect of simplified model parameters on the response to periodic stimuli (***Figure 5—figure supplement 3***), using the same parameter variations explored for pulse stimuli. The state duration scales could dramatically alter the entrainment response, which arose from relocated or removed stable points in the phase response curve. The adaptation time constant and scales increased entrainment due to the larger and more stable phase shift response. The quiet switching delay increased entrainment for longer-period, inhibitory pulse trains due to the wider stabilizing region in the phase response curve. These results suggest parameters to tune for a desired entrainment response.

### Simplified model can be integrated into neural circuits

We evaluated the simplified model’s advantages for circuit modeling by integrating it into 2 neural circuit models—the crustacean pyloric circuit and the mammalian spinal locomotor circuit. These circuits are well-studied and have been simulated using different types of neuron models, including detailed conductance-based spiking models and reduced non-spiking models. We reproduced key circuit activity patterns from reference works using the simplified model.

#### Crustacean pyloric circuit generates a triphasic rhythm

The pyloric circuit is a CPG located in the stomatogastric ganglion of crustaceans that controls the pyloric muscles of the stomach (***Marder and Bucher, 2007; Maynard, 1972***). It is composed of an electrically coupled pacemaking kernel of anterior burster (AB) and pyloric dilator (PD) neurons, connected through inhibitory synapses to lateral pyloric (LP) and pyloric (PY) neurons that are mutually inhibiting. The circuit generates a periodic, triphasic rhythm during which intrinsically bursting AB neurons synchronously drive PD neurons to burst, followed by LP neurons then PY neurons that are activated through post-inhibitory rebound operating with different delays. For details, see ***Marder and Bucher (2007)***.

We used as reference the work of ***Alonso and Marder (2020)***, which modeled the pyloric circuit using detailed conductance-based spiking neurons within a simplified architecture of 3 neurons (***Figure 6A***). Following ***Liu et al. (1998)***, each neuron was a dynamical system of 13 state variables: 1 voltage variable, 1 calcium variable, and 11 activation or inactivation gating variables used by a total of 8 currents—sodium (Na), transient calcium (CaT), slow calcium (CaS), transient potassium (A), calcium-dependent potassium (KCa), delayed rectifier potassium (Kd), hyperpolarization-activated inward (H), and leak (L). Each current required parameters for its maximal conductance and reversal potential. Each gating variable required parameters for its exponent, as well as its voltage- or calcium-dependent steady-state and time constant functions. Additionally, the model incorporated chemical synapses with conductance-based dynamics of various timescales (adding 7 state variables), as well as temperature-dependent scaling for the 8 maximal conductances, 11 gating variable time constants, 1 calcium time constant, and 4 synaptic parameters (adding 24 parameters). The model parameters were tuned using an multi-stage evolutionary search algorithm (***Alonso and Marder, 2019***) on objective functions that quantified desired characteristics for the rhythm, such as burst frequencies, duty cycles, and phase lags. Through optimization, a temperature-robust model was found that generated stable duty cycle and phase relationships.

**Figure 6.**
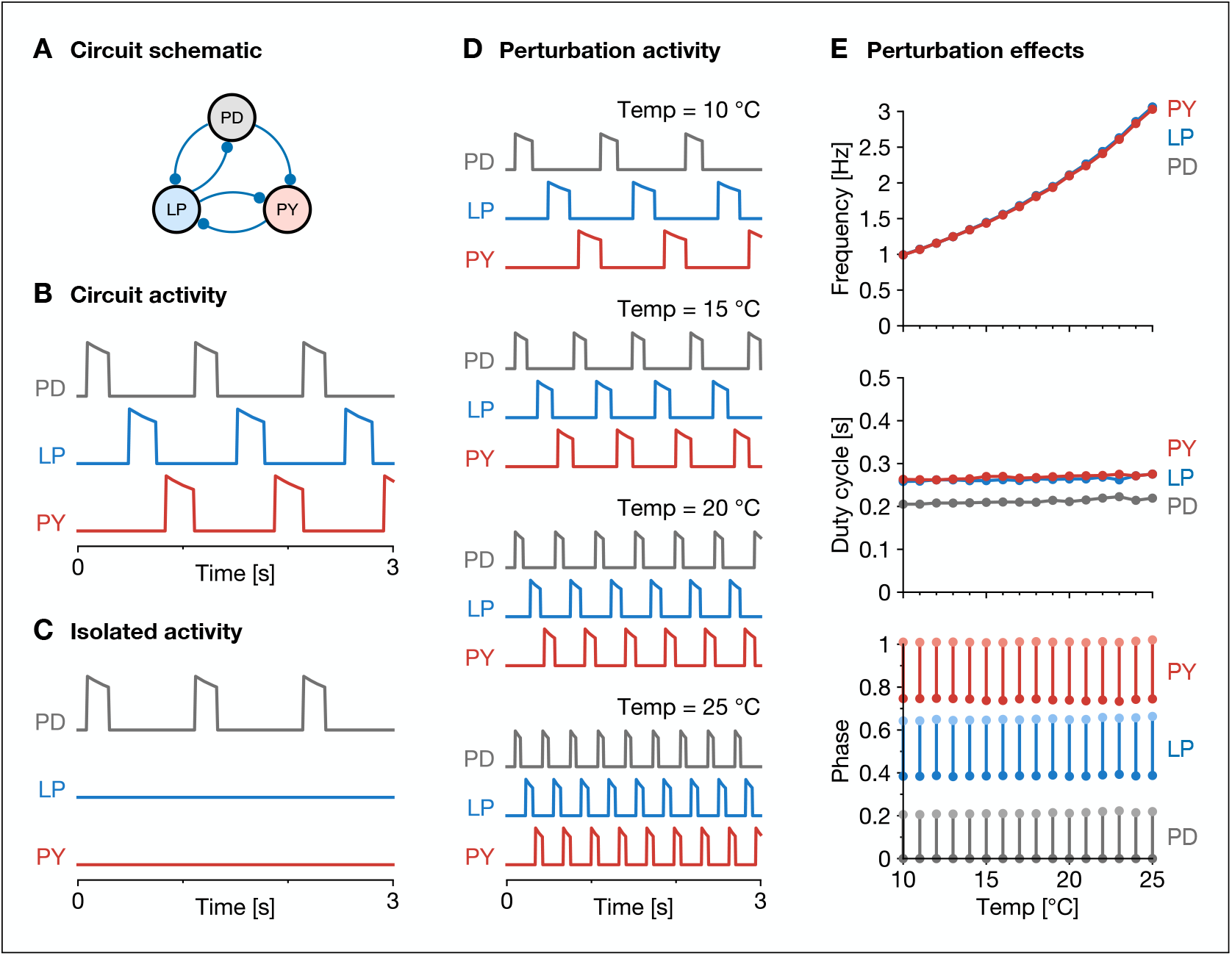
Crustacean pyloric circuit. (**A**) Schematic of the circuit adapted from ***Alonso and Marder (2020)***. The 3 neurons are connected through inhibitory synapses (blue circles). (**B**) Activity of the circuit with intact synapses. A periodic, triphasic rhythm emerges in which neurons burst in sequence: PD → LP → PY. (**C**) Activity of the circuit with isolated neurons after ablating synapses. Only the PD neuron bursts intrinsically, indicating that the LP and PY neurons were activated by post-inhibitory rebound effects. (**D**) Activity of the circuit under temperature perturbation, which was modeled as scaling of time-based parameters. (**E**) Effects of temperature perturbations on each neuron’s frequency, duty cycle, and phase. The simplified model reproduces these phenomena from ***Alonso and Marder (2020)***, in which burst frequency increases with temperature, but the average duty cycle and phase relationships between neurons remain constant.

With the simplified model, we reproduced key circuit activity patterns and temperature perturbation effects from ***Alonso and Marder (2020)***. Due to the simplified model’s interpretable and controllable parameters, we could easily select parameters to match experimental observations (***Table 3***). Specifically, active and quiet durations of neurons were selected to establish desired burst frequencies and duty cycles, while quiet switching delays of LP and PY were selected to establish desired phase relationships. The circuit produced the characteristic triphasic rhythm, with sequential bursting of PD then LP then PY (***Figure 6B***). Moreover, ablating synapses to isolate the neurons demonstrated that only PD continued to burst intrinsically, while LP and PY were silent (***Figure 6C***). This reflected how LP and PY were subthreshold at baseline input and were activated in the intact circuit through post-inhibitory rebound effects, consistent with experimental observations (***Marder and Bucher, 2007***). Finally, temperature dependence was introduced as a shared scaling term on time-based parameters, which comprised the state durations, adaptation time constant, and stable point time margins. This enabled burst frequency to increase with temperature (***Figure 6D***), while keeping constant the duty cycle and phase relationships between neurons (***Figure 6E***). These results were qualitatively and quantitatively similar to Figure 1 of ***Alonso and Marder (2020)*** and may in fact score better on the optimization objectives, since the simplified model enabled the duty cycle and phase relationships to remain exactly at desired values, unlike the conductance-based model that exhibited some variability due to its mechanisms of current compensation.

**Table 3.** Simplified model parameters for circuit experiments.

| Symbol | Pyloric |  |  | Locomotor |  |
| --- | --- | --- | --- | --- | --- |
|  | PD | LP | PY | Flx | Ext |
| $\Delta t$ | 1 ms | 1 ms | 1 ms | 1 ms | 1 ms |
| $T_{\text{active}}$ | 200 ms | 250 ms | 250 ms | 75 ms | 75 ms |
| $T_{\text{quiet}}$ | 800 ms | 750 ms | 750 ms | 400 ms | 400 ms |
| $K_{\text{active}}^{+1}$ | 1.0 | 1.0 | 1.0 | 0.125 | 0.125 |
| $K_{\text{quiet}}^{+1}$ | 1.0 | 1.0 | 1.0 | 0.025 | 0.025 |
| $K_{\text{active}}^{-1}$ | 0.9 | 0.5 | 0.5 | 0.4 | 0.4 |
| $K_{\text{quiet}}^{-1}$ | 1.1 | 1.3 | 1.3 | 1.1 | 1.1 |
| $T_a$ | 1500 ms | 1500 ms | 1500 ms | 450 ms | 450 ms |
| $K_{a,\text{active}}^{+1}$ | 1.0 | 1.0 | 1.0 | 1.0 | 1.0 |
| $K_{a,\text{quiet}}^{+1}$ | 1.0 | 1.1 | 1.1 | 1.2 | 1.2 |
| $K_{a,\text{active}}^{-1}$ | 1.0 | 1.1 | 1.1 | 1.2 | 1.2 |
| $K_{a,\text{quiet}}^{-1}$ | 1.0 | 0.5 | 0.5 | 0.8 | 0.8 |
| $D_{\text{active}}$ | 0.0 | 0.0 | 0.0 | 0.0 | 0.0 |
| $D_{\text{quiet}}$ | 0.0 | 0.25 | 0.95 | 0.1 | 0.1 |
| $X_{\text{active}}$ | 1.0 | 1.0 | 1.0 | 1.0 | 1.0 |
| $X_{\text{quiet}}$ | 0.0 | 0.0 | 0.0 | 0.0 | 0.0 |
| $\delta_{\text{active}}$ | 0 ms | 0 ms | 0 ms | 0 ms | 0 ms |
| $\delta_{\text{quiet}}$ | 10 ms | 10 ms | 10 ms | 50 ms | 50 ms |
| $B$ | 0.0 | -0.1 | -0.1 | -0.05 | 1.2 |
| $\sigma$ | 0.05 | 0.05 | 0.05 | 0.02 | 0.02 |
| $y$ | $y_{\text{adapting}}$ | $y_{\text{adapting}}$ | $y_{\text{adapting}}$ | $y_{\text{curved}}$ | $y_{\text{curved}}$ |

#### Mammalian locomotor circuit generates gait patterns

The locomotor circuit is a CPG located in the spinal cord of mammals that produces gait rhythms for quadruped locomotion (***Rybak et al., 2015***). It is composed of reciprocally inhibiting flexor and extensor bursting half-centers for each limb, which are connected through excitatory and inhibitory synapses to non-bursting spinal interneurons. Brainstem command (BC) connections provide top-down modulation to half-centers and interneurons, controlling burst frequencies and limb synchonization or alternation. The circuit generates periodic rhythms during which limb half-centers burst to produce fictive gait patterns. As BC increases, the gait transitions from stand to walk to trot to gallop to bound. For details, see ***Rybak et al. (2015); Danner et al. (2016)***.

We used as reference the work of ***Danner et al. (2017)***, which modeled the locomotor circuit using reduced non-spiking neurons corresponding to the biophysical model studied in the present work. The circuit used a simplified architecture in which single neurons represented populations of genetically defined neuron types. The original architecture has since been extended by ***Ausborn et al. (2019)*** and ***Zhang et al. (2022)*** with new connections, which we incorporated into our simplified architecture (***Figure 7A***). In ***Danner et al. (2017)***, non-bursting neurons were modeled by omitting the persistent sodium current (NaP), and synapses were modeled without dynamics. The model parameters were manually tuned to achieve desired locomotor frequencies and gaits.

**Figure 7.**
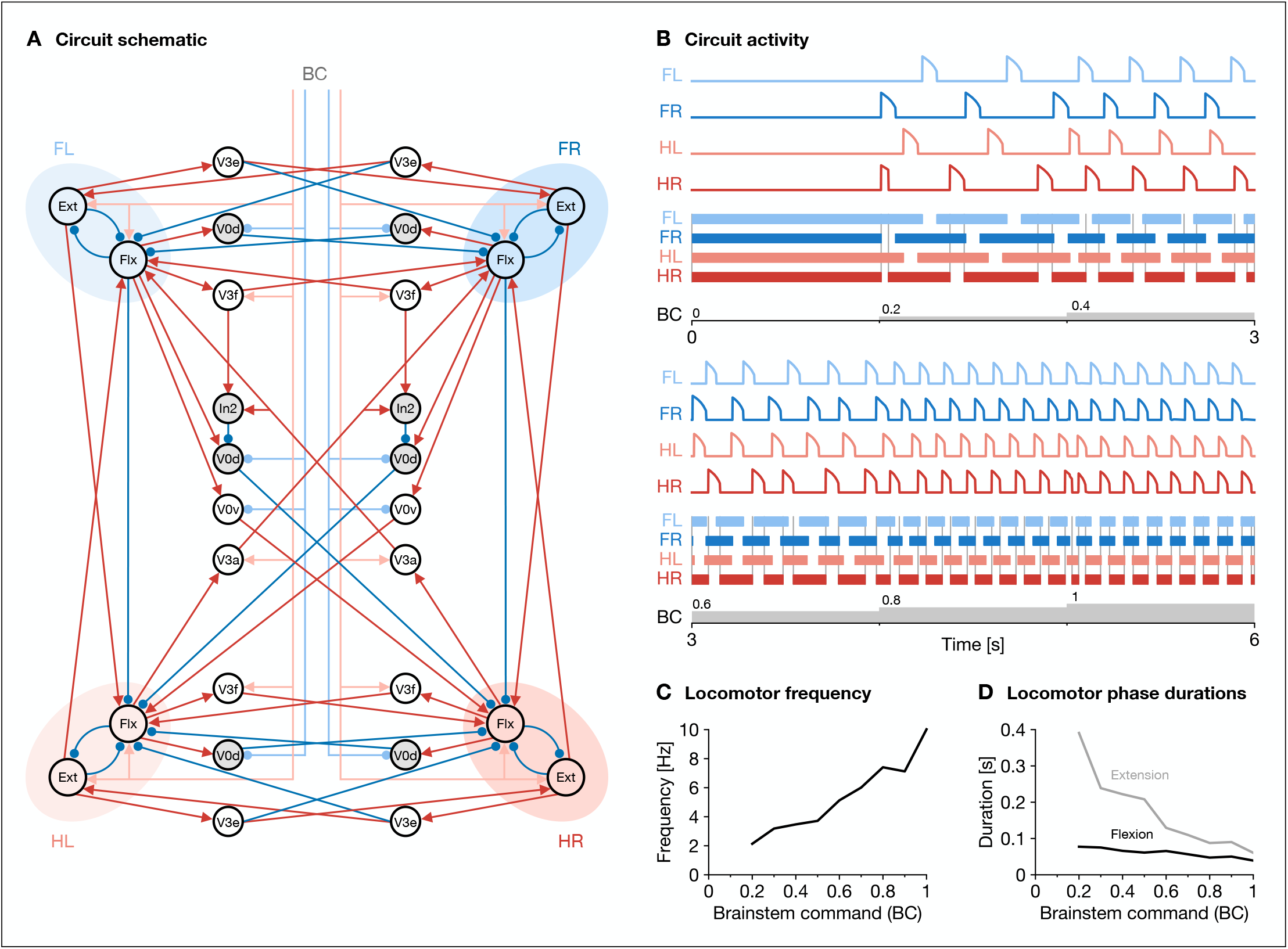
Mammalian locomotor circuit. (**A**) Schematic of the circuit adapted from ***Danner et al.(2017); Ausborn et al. (2019); Zhang et al. (2022)***. The 4 limbs are each controlled by reciprocally inhibiting flexor and extensor bursting half-centers, which are connected through excitatory synapses (red arrows) and inhibitory synapses (blue circles) to non-bursting spinal interneurons. Brainstem command (BC) provides top-down modulation to half-centers and interneurons. Limbs are named fore-left (FL), fore-right (FR), hind-left (HL), and hind-right (HR). (**B**) Activity of the circuit in response to increasing brainstem command (BC). Top: Flexor activity for each limb. Bottom: Extensor activity for each limb (rendered as a simulated footfall plot to facilitate gait analysis). As BC increases, the gait transitions from: stand → walk → trot → gallop → bound. (**C**) Locomotor frequency increases with increasing BC. (**D**) Locomotor phase durations decrease with increasing BC, with the extension duration decreasing sharply compared to the flexion duration. The simplified model reproduces these phenomena from ***Danner et al. (2017***). **Figure 7—figure supplement 1**. Activity of neuron types at different locomotor frequencies.

With the simplified model, we reproduced key circuit activity patterns and locomotor effects from ***Danner et al. (2017)***. We selected state durations and scales to establish desired burst frequencies and duty cycles at boundary BC values, and we tuned neuronal biases and synaptic weights to produce the appropriate gaits across BC values. The circuit produced flexor and extensor activity that transitioned between gaits as BC increased (***Figure 7B***), including a stable stand gait at low BC not shown in the reference work. These gaits were reflective of BC-dependent activation of various interneurons types (***Figure 7—figure supplement 1***), with interneuron activity closely matching ***Zhang et al. (2022)***, which used detailed conductance-based spiking neurons. Furthermore, the model reproduced key metrics of the locomotor rhythm. As BC increased from 0 to 1, locomotor frequency increased from 2 Hz to 10 Hz (***Figure 7C***), and phase durations both decreased, with extension decreasing more sharply than flexion (***Figure 7D***). These results were qualitatively and quantitatively similar to Figure 4 from ***Danner et al. (2017)***, which in turn conformed to experimental observations of mouse locomotion.

## Discussion

We sought a simplified model of intrinsically bursting neurons that was computationally efficient, easily tunable, and readily integrated into neural circuits. Our model achieves these goals while reproducing single-unit responses and circuit-level activity patterns of various biophysical models.

Our model’s core mechanism involves analytically computing the active and quiet bounds to achieve desired state durations and input-dependent scaling (***Equation 20, Equation 24***). This is possible since adaptation *a* is governed by a piecewise-linear and separable differential equation. Prior work has explored piecewise-linear approximations, notably the McKean model (***McKean, 1970***), which used a piecewise-linear “Z”-shaped function to approximate the voltage nullcline of the FitzHugh-Nagumo model (***FitzHugh, 1961; Nagumo et al., 1962***) while maintaining the latter’s continuous dynamics and biphasic truncated exponential waveform. In contrast, our model uses a piecewise-linear “S”-shaped function consisting of horizontal and vertical segments, which decouples movement of the voltage and adaptation variables (***Figure 1C***). Our design has the advantage of facilitating precise movement control on horizontal segments to scale state durations as well as on vertical segments to produce state switching delays. Moreover, having trajectories move along a rectangular limit cycle facilitates transforming state variables into firing activity waveforms with different burst shapes (***Figure 2C***). However, our design incurs the cost of using hybrid continuous-discrete rules to govern dynamics, which introduces complexity and requires explicit handling of dynamical properties such as stable points. Nevertheless, this tradeoff seems acceptable since it affords additional control of those dynamical properties, and since discrete rules are already commonly used in simplified models (***Izhikevich, 2003***).

In certain respects, our model is reminiscent of phase oscillator models. It focuses on capturing the core limit cycle of bursting dynamics using state variables that oscillate in an abstract coordinate system (***Buchli et al., 2006***), and it decouples dynamics that are tangential or perpendicular to the limit cycle. It also relies on a transformation—the firing function—to convert between state variables and biologically meaningful quantities. However, our model explicitly represents a rectangular limit cycle in Cartesian coordinates, reflecting its roots in the biophysical model. Thus, state variables and phase portraits closely correspond with biophysical interpretations, and it is easier to incorporate biophysically relevant dynamical properties such as stable point rebound, which depend on representations of input-dependent active and quiet bounds.

Despite its similarities to the biophysical model, our model’s approximation has limitations. First, the voltage variable is an abstract quantity that oscillates between –1 and +1, whereas the biophysical model’s voltage range is biologically meaningful and can shift with input, even shifting above threshold entirely at strong excitatory input (***Figure 3—figure supplement 1***). This discrepancy would be relevant for features like gap junctions where voltage differences matter, so some function would be needed to transform the variable, similar to the firing function. Second, the voltage dynamics produce instantaneous jumps from the threshold when switching states (***Equation 27***), which simplifies calculations but leads to sharp burst onsets. Smooth onsets for burst shapes like sinusoids would require modifying the firing function or introducing graded movement similar to state switching delay. Third, the adaptation dynamics uses a single time constant for depletion and recovery (***Equation 19***). This could easily be relaxed to allow asymmetric rates using an additional parameter that makes one a fixed multiple of the other. Fourth, the adaptation variable monotonically depletes or recovers except when instantaneously clipped to bounds around stable points (***Equation 30***). This approximation is only realistic when time margins are no greater than the discretization timestep; otherwise, the trajectory should take multiple timesteps to return to the stable point with reversed movement, though in practice this edge case has not been an issue.

Compared to detailed conductance-based models, our model—like other reduced non-spiking models—does not reproduce the full richness of burst spiking dynamics, especially in extreme conditions. For example, ***Alonso and Marder (2020***) show that neurons can “crash” when ion channel kinetics are modulated by extreme perturbations, producing irregular and aperiodic spiking activity. In contrast, our model is well-behaved even at extreme inputs, which may be desirable for constructing robust circuits for neuro-robotics but is nonetheless biologically unrealistic.

Yet in some cases, our model could produce more biologically realistic activity than the biophysical model. For example, ***Shevtsova et al. (2015)*** simulated a half-center microcircuit using detailed conductance-based spiking neurons. As input increased, activity transitioned from silent to bursting to tonic mode, with burst amplitude slowly ramping up then down, peaking midway through bursting mode. However, when the microcircuit was adapted by shared authors in ***Danner et al. (2017)*** to use the biophysical model, this amplitude modulation response was not maintained. Instead, amplitude immediately jumped to its peak at the silent-to-bursting transition then ramped down (***Figure 3—figure supplement 1***). It is unclear how to adjust the biophysical model to reproduce the response, since it involves altering ion channel parameters that shape the voltage nullcline. With our simplified model, such amplitude modulation is straightforward (***Figure 1B***).

Beyond producing activity, the simplified model offers insight into how specific responses arise in the biophysical model. By abstracting active and quiet bound shifts, it becomes evident how oscillations in different regions of the asymmetric adaptation field influence the duty cycle of the bursting waveform (***Figure 3—figure supplement 2***). Moreover, by decoupling bound shifts and adaptation time constant scaling, the model reveals how each mechanism distinctly shapes the phase response curve (***Figure 4—figure supplement 3***). These relationships, complicated by non-linear interactions in the biophysical model, became readily interpretable in the simplified model.

Applications of the simplified model can leverage its interpretability and controllability. The model can be used to rapidly prototype neural circuits to match experimental observations. It can also complement biologically detailed models by serving as a surrogate during optimization, producing activity targets that guide parameter tuning. Additionally, the model can be used to synthesize novel CPG architectures with desired activity patterns. Notably, we have validated its applicability to neuro-robotics, where it served as a core component of a neural architecture controlling quadruped locomotion on a physical robot (***Bhattasali et al., 2024***).

Future work can extend the simplified model in several directions. It could be integrated into more complex neural circuits, including those involving gap junctions or neuromodulatory influences. It could also be augmented with intrinsic plasticity mechanisms (***Righetti et al., 2006***), enabling parameters to update based on long-term activity patterns. These extensions would expand the simplified model’s utility for biological circuit modeling and neuro-robotics.

## Methods

### Biophysical model

#### Equations

The biophysical model is based on the reduced non-spiking neuron from ***Danner et al. (2017***) described as a 2-dimensional dynamical system with voltage variable *v* and adaptation variable *a*.

The voltage variable *v* ∈ ℝ represents the neuron’s average membrane potential governed by a differential equation relating membrane capacitance *C* and various trans-membrane ionic currents *i*, each with its own maximal conductance *G* and reversal potential *V* :

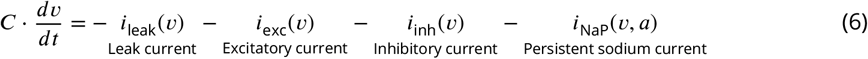

The leak current maintains the resting membrane potential:

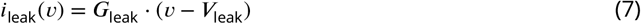

The excitatory and inhibitory synaptic currents depolarize and hyperpolarize, respectively, the membrane potential in response to constant drives *D* and synaptic input, summed over presynaptic neurons *n* with synaptic weights *w*_*n*_ ≥ 0 and presynaptic firing activities *y*(*v*_*n*_) ≥ 0:

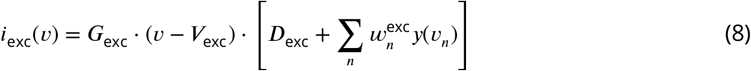

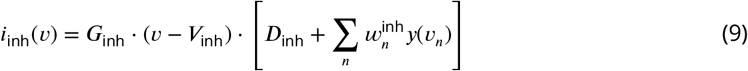

The persistent sodium (NaP) current produces bursting through its slow dynamics:

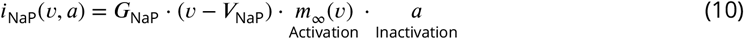

where activation is modeled as instant with the steady-state activation *m*_∞_(*v*):

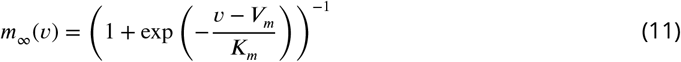

and inactivation is modeled as slow with the adaptation variable *a* ∈ [0, 1] governed by a differential equation relating the steady-state inactivation *a*_∞_(*v*) and time constant τ_*a*_(*v*):

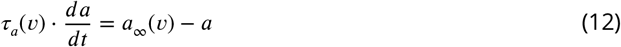

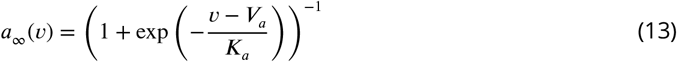

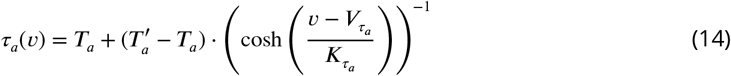

and various parameters (*V, K, T*) control the shape of sigmoid and bell curves.

The firing activity of the neuron is a non-negative function of the voltage variable *v*:

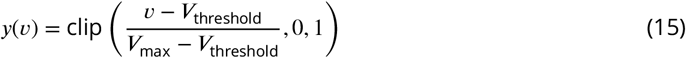

#### Discretization

For simulation, the differential equations for voltage (***Equation 6***) and adaptation (***Equation 12***) were discretized and integrated using the Forward-Euler method due to its simplicity with inseparable nonlinear terms:

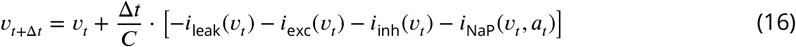

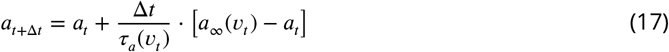

### Simplified model

#### Equations: Bounds

The simplified model is built around a core mechanism for controlling active and quiet durations by setting appropriate bounds for the active and quiet nullcline legs. This section derives the core mechanism by considering the simplest case, with fixed adaptation time constant and state durations (no input modulation) and instantaneous state switches between legs at the bounds (no delay, *v* ∈ {−1, +1}). The following sections elaborate the core mechanism with additional features.

Formally, given the adaptation time constant *T*_*a*_ (controlling the rate of movement along the legs) and the active duration *T*_active_ and quiet duration *T*_quiet_ (specifying the desired time spent on each leg), the goal is to determine the active bound *a*_active_ and quiet bound *a*_quiet_ that together trigger state switches at the desired times.

The adaptation variable *a* ∈ (0, 1) depletes toward 0 (“empty”) in the active state and recovers toward 1 (“full”) in the quiet state, following an exponential trajectory that approaches, but never reaches, those asymptotic values:

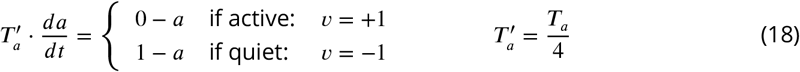

For convenience, the adaptation time constant *T*_*a*_ is made more interpretable through scaling: Since an exponential trajectory reaches 98.2% ≈ 1 − exp(−4) toward its asymptote after 4 time constants, 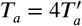 can be interpreted as the time it takes for the adaptation variable to deplete or recover “almost completely” when starting from full or empty, respectively.

The time spent on each leg is analytically derived by integrating the time-per-unit-adaptation between the (unknown) bounds. The time-per-unit-adaptation is the inverse of the adaptation-per-unit-time, which is the adaptation rate in ***Equation 18***:

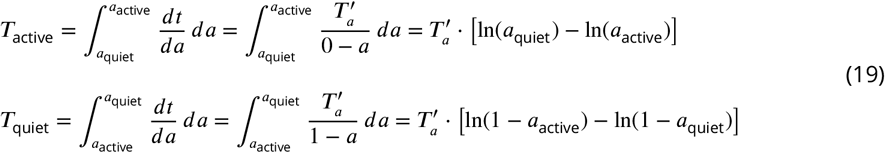

This yields a system of 2 equations with 2 unknowns, which can easily be solved for the bounds by applying logarithmic properties, exponentiation, and substitution:

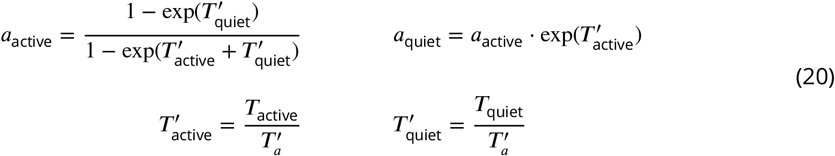

Thus, these bounds *a*_active_ and *a*_quiet_ trigger state switches at the desired times.

#### Equations: Modulation

The simplified model can modulate its properties in response to net input *x*, which includes bias input *B* and synaptic input, summed over presynaptic neurons *n* with synaptic weights *w*_*n*_ ∈ ℝ and presynaptic firing activities *y*_*n*_ ≥ 0:

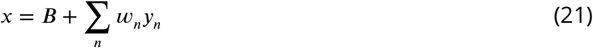

Specifically, the net input scales the adaptation time constant *T*_*a*_ (using 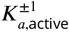 and 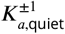) and the state durations *T*_active_ and *T*_quiet_ (using 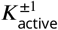 and 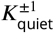).

The adaptation variable differential equation from ***Equation 18*** is modified to use 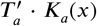 *K*_*a*_(*x*), incorporating an input-dependent scaling factor for the adaptation time constant:

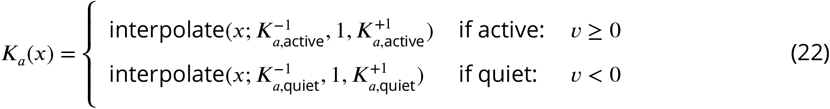

which applies a two-slope linear interpolation over *x* (clipped between −1 and +1) to interpolate between the left (*x* = −1), middle (*x* = 0), and right (*x* = +1) values.

The active and quiet bounds are also modified to be input-dependent by calculating the bounds from ***Equation 20*** for no input (*x* = 0) and the strongest inputs (*x* = ±1):

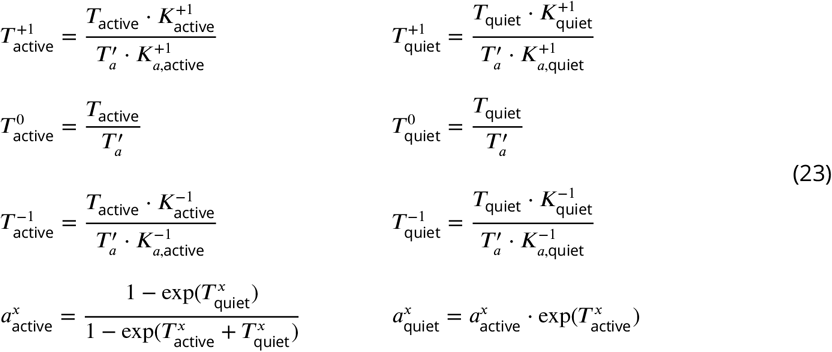

and then linearly interpolating between the bounds for intermediate inputs:

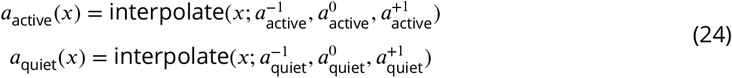

Thus, desired state durations are achieved even as the adaptation time constant is modulated.

#### Equations: Delay

The simplified model can switch between states instantaneously or with some delay. Such delay can be useful for establishing phase relations between neurons (***Figure 6***) or enabling a period of competition between neurons during which state switches can be rejected (***Figure 7***). Specifically, delay is specified as a fixed fraction *D*_active_, *D*_quiet_ ∈ [0, 1] of the input-dependent state durations:

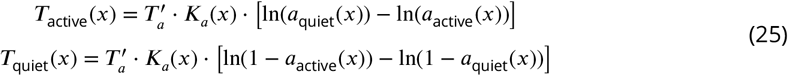

Without delays, a trajectory moves along only horizontal segments of the active and quiet legs (***Figure 1C***) then instantaneously switches legs at the bounds. With delays, a trajectory moves along the horizontal segments then turns at the bounds to move along the vertical segments, only switching legs once *v* crosses zero. Across these cases, the time spent on each leg must be conserved in order to achieve the desired state durations. Therefore, delays can intuitively be interpreted as “stealing time” from horizontal segments and shifting it to vertical segments.

The adaptation variable differential equation from ***Equation 18*** and ***Equation 22*** is modified to speed up movement along horizontal segments and stop movement outside bounds:

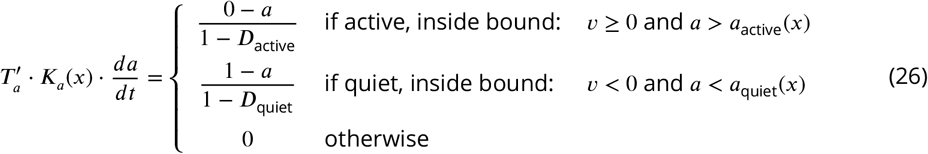

The voltage variable differential equation is defined to introduce movement at a constant rate along vertical segments and switch legs at zero:

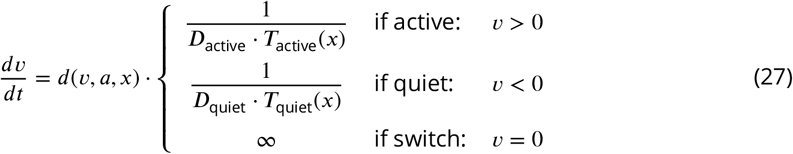

where the direction function *d*(*v, a, x*) specifies the direction of movement within each leg region using sign(*z*) : ℝ → {−1, 0, +1}:

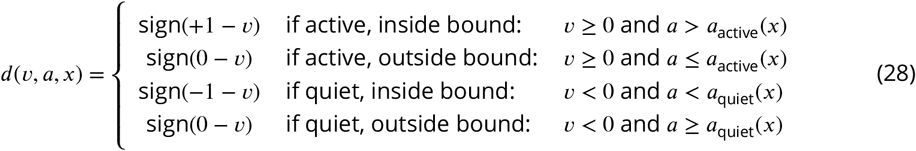

and instantaneous switches (of infinite voltage rate due to *v* = 0 or any *D* = 0) are handled by continuously constraining *v* to the range [−1, +1].

Thus, instantaneous and delayed state switches are achieved with unified dynamics.

#### Equations: Stability

The simplified model can enter stable modes in which bursting is suppressed. Specifically, it enters the tonic mode at net input above the active stable point threshold *X*_active_ ≥ 0, and it enters the silent mode at net input below the quiet stable point threshold *X*_quiet_ ≤ 0.

In stable modes, a trajectory moves along the active and quiet legs but stops at the corner point on the stable bound, without moving onto the subsequent vertical segment. The trajectory is only released once net input no longer meets threshold. If a trajectory is already moving vertically when net input meets threshold, then it reverses direction and returns to the stable point. Importantly, the trajectory only reverses if it is close enough horizontally to the bound (in terms of adaptation); otherwise, it continues vertically to trigger a state switch known as a rebound, which is non-periodic and terminates when the trajectory returns to the stable point after a single burst.

The voltage variable direction function is modified to reverse the direction of movement along vertical segments in response to net input:

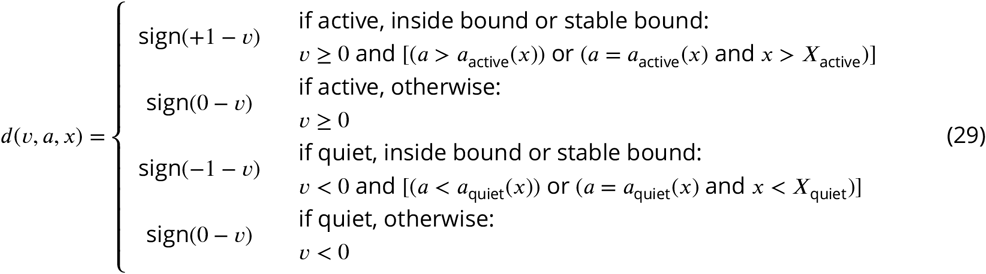

The adaptation variable dynamics is modified to include an instantaneous constraint that clips adaptation to bounds if inside bound margins *ã*_active_(*x*), *ã*_quiet_(*x*):

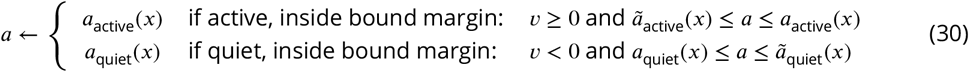

where the bound margins are calculated from time margins *δ*_active_, *δ*_quiet_ similar to in ***Equation 19*** and ***Equation 20*** (derivation omitted):

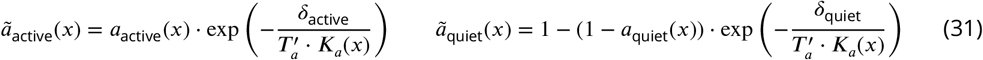

Thus, stable modes with sophisticated dynamics are expressed in a highly controllable manner.

#### Equations: Noise

The simplified model can incorporate noise into the dynamics, which is useful for increasing variability and breaking symmetries. Specifically, it scales voltage and adaptation rates with non-negative multiplicative noise *ε* sampled at each timestep from a normal distribution with mean 1 and standard deviation *σ*:

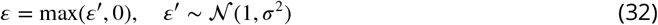

Thus, state durations are sped up or slowed down while maintaining their relative timing.

#### Equations: Activity

The simplified model can decouple the shape of active and quiet legs from the activity waveform’s burst amplitude and shape through the firing function *y*(*v, a, x*), which produces non-negative firing rate activity based on the neuron’s state and net input (***Figure 2***).

A simple *rectangular waveform* can use constant values along both active and quiet legs:

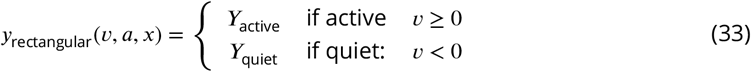

or an *adapting waveform* can linearly interpolate between high and low values along the active leg:

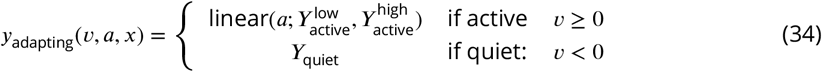

where linear interpolation between endpoint values *A, B* uses *z* ∈ [0, 1]:

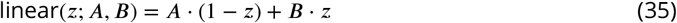

A more controllable *curved waveform* can set exact values at the bounds and adjust the curve to be convex, concave, or approximately linear (by adjusting *C*):

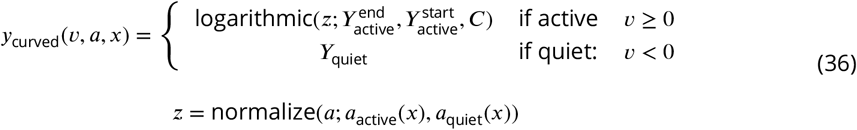

where logarithmic interpolation between endpoint values *A, B* uses *z* ∈ [0, 1] and base *C* > 0:

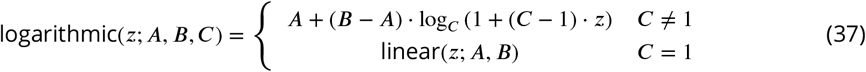

and bounds normalization between bound locations *A, B* uses *u* ∈ ℝ:

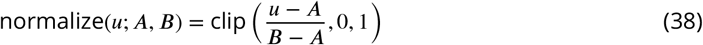

Thus, a variety of waveform shapes are achieved with the same underlying dynamics.

#### Discretization

For simulation, the differential equations for voltage (***Equation 27***) and adaptation (***Equation 26***) were discretized and integrated using the Backward-Euler method due to its stability and tractability with separable piecewise-linear terms. The order of integration and constraint application is important to maintain the correct logic.

First, the current adaptation is clipped to the new bounds if within time margins (***Equation 30***). Second, the new adaptation is calculated from the dynamics, using a newly sampled noise:

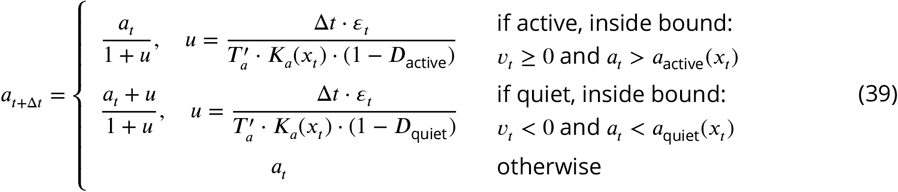

Third, the new adaptation is clipped to bounds if needed to fix overshoot within the timestep.

Fourth, the new voltage is calculated from the dynamics, using the new adaptation to move in the correct direction along the bounds:

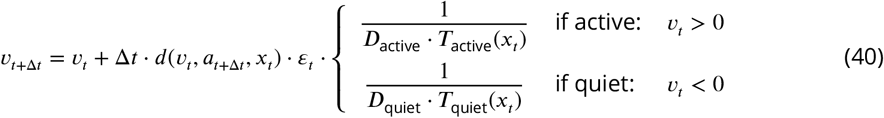

Fifth, the new voltage is set to +1 or –1 to implement an instantaneous state switch if it crossed zero within the timestep.

### Simulation software

Models were simulated and analyzed in Python on Linux machines with 16GB of shared RAM memory. Computations were parallelized across 8 to 16 CPU workers to ensure completion in under 30 minutes. A Miniconda virtual environment was used to manage dependencies. Code is open-sourced on GitHub (https://github.com/nikhilxb/bursting) to facilitate reproducibility.

### Single-unit experiments

#### Parameters

Single-unit experiments were run using a consistent set of parameters, and an additional parameter *I* was introduced to net input *x* to represent stimulus input. For the biophysical model, parameters were used from ***Kim et al. (2022***), which hand-tuned them to match experimental observations of cat locomotion (***Table 1***). For the simplified model, parameters were hand-tuned to match the biophysical model in response to constant, pulse, and periodic stimuli (***Table 2***). The firing function used a customized *y*_curved_ to approximate the biophysical model’s input-dependent changes to nullcline-*v* (and therefore the shape of its activity waveform):

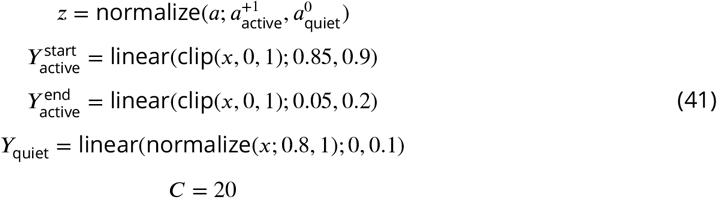

#### Waveform analysis

A waveform’s cycle duration, along with its active and quiet durations, were calculated from zero-crossing times, which are when the waveform rises above or falls below zero. An active phase is bounded by a rise and the subsequent fall, while a quiet phase is bounded by a fall and the subsequent rise. A waveform’s peaks were identified using a SciPy function that detects local maxima in a 1D sequence of values by comparing each value to its neighbors.

#### Constant stimuli analysis

Constant stimuli of various strengths were applied to assess duration scaling. For each response waveform, the intrinsic cycle duration was measured, along with the active and quiet durations. Duty cycle was computed as the ratio of active duration to intrinsic cycle duration. These measurements were limited to waveforms with both active and quiet phases, excluding cases where the model was in tonic/silent mode or produced entirely non-zero firing activity.

#### Pulse stimuli analysis

Pulse stimuli of various strengths and phases were applied to assess phase shifts. Pulses were 100 ms in duration. At each strength, pulses were tested across the intrinsic cycle at pulse start times separated by 25 ms. Phase shift Δ*ϕ* was quantified by comparing the perturbed cycle duration *T* ^′^ to the intrinsic cycle duration *T* :

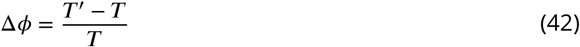

#### Periodic stimuli analysis

Periodic stimuli of various strengths and periods were applied to assess entrainment. Pulse trains were tested with normalized periods (relative to the intrinsic cycle duration) from 0.75 to 1.25 in steps of 0.05. Pulses were 100 ms in duration. Pulse trains contained 10 pulses, chosen to balance the tradeoff between capturing transient and steady-state effects. At each strength and period, pulse trains were tested with initial pulse phases from 0 to 1 in steps of 0.1.

Entrainment was quantified by phase coherence. For each response waveform, burst peaks were identified and assigned phases *ϕ* ∈ [0, 1) relative to the stimulus. These phases converge to a steady-state value under entrainment, and their circular mean characterizes convergence speed and robustness. The circular mean phase 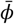 was calculated across pulses and initial pulse phases:

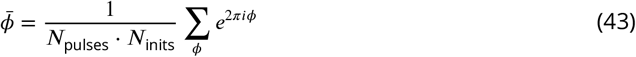

Phase coherence is the magnitude of this complex-valued 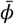, with a value of 1 if all burst peaks align at the same phase relative to the stimulus. In practice, phase coherence is less than 1 even for fast and robust entrainment due to initial transient effects.

### Circuit experiments

#### Equations: Basic

Circuits use basic neurons in addition to bursting neurons. The simplified model of basic neurons used for experiments has standard linear dynamics that can modulate firing rate in response to net input *x*, which includes bias input *B* and synaptic input, summed over presynaptic neurons *n* with synaptic weights *w*_*n*_ ∈ ℝ and presynaptic firing activities *y*_*n*_ ≥ 0:

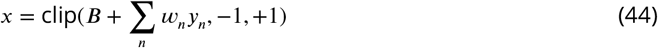

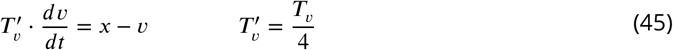

For convenience, the voltage time constant *T*_*v*_ is made more interpretable through scaling, as in ***Equation 18***.

The firing activity of the neuron is a non-negative function of the voltage variable *v*:

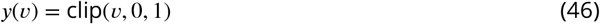

For simulation, the differential equation for voltage was discretized and integrated using the Backward-Euler method due to its stability and tractability with linear terms:

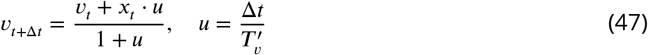

#### Parameters

Circuit models included parameters for neuronal dynamics and synaptic weights. For bursting neurons, the parameters were specific to each circuit (***Table 3***). For basic neurons, the parameter *T*_*v*_ = 10 m*s* was used for all circuits. Biases and synaptic weights were hand-tuned to reproduce the circuit activity in reference works. The full list of these parameter values can be found in the code.

For the crustacean pyloric circuit, the triphasic rhythm was targeted to a period of 1 second, with PD active for 200 ms, LP for 250 ms, and PY for 250 ms. Quiet switching delays of LP and PY established the appropriate firing phases following post-inhibitory rebound induced by PD. Temperature *c* (in Celsius) was incorporated as a scaling factor *τ*(*c*) on the adaptation time constant *T*_*a*_, active duration *T*_active_, quiet duration *T*_quiet_, and quiet stable point time margin *δ*_quiet_:

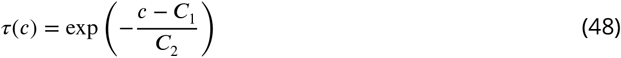

with constants *C*_1_ = 10 ^◦^C and *C*_2_ = 13 ^◦^C to match the empirically observed scaling relation. The firing function used *y*_adapting_ with 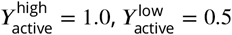, and *Y*_quiet_ = 0.

For the mammalian locomotor circuit, flexor and extensor centers used the same set of parameters except biases, which were set differently to ensure that extensor centers operated in tonic mode at baseline input while flexor centers operated in bursting mode. Synaptic weights were shared bilaterally to exploit symmetry. The firing function used *y*_curved_ from ***Equation 41*** with *Y*_quiet_ modified to begin rising at net input of 0.6 (rather than 0.8).

## Acknowledgments

Thanks for insightful discussions to Pavel Tolmachev. This work was supported by the Fannie and John Hertz Foundation Fellowship (NXB); NSF award 2339096 (LP); ONR awards N00014-21-1-2758 and N00014-22-1-2773 (LP); and the Packard Fellowship for Science and Engineering (LP).

---

**Figure 3—figure supplement 1.**
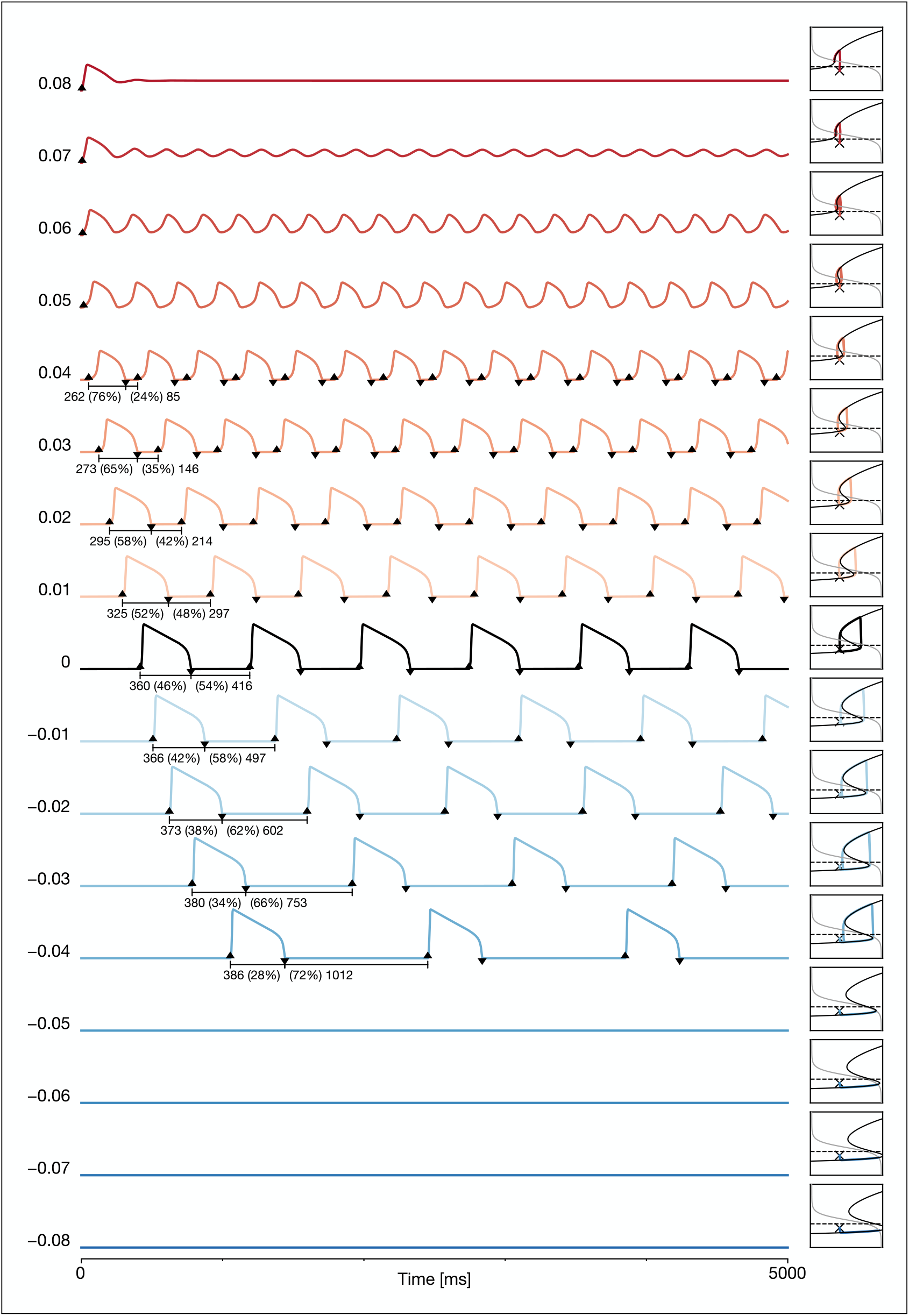
Biophysical model response waveforms to constant stimuli. Activity for all stimulus strengths shown in ***Figure 3***, along with the corresponding phase portraits. For stimulus strengths above 0.04, the activity does not have a quiet phase as the neuron fires at low rates between bursts. For stimulus strengths above 0.08, the neuron enters a stable tonic firing mode. For stimulus strengths below –0.04, the neuron enters a stable silent mode. Intervals show the active and quiet durations (in milliseconds) and the duty cycle (as a percentage of the oscillation period).

**Figure 3—figure supplement 2.**
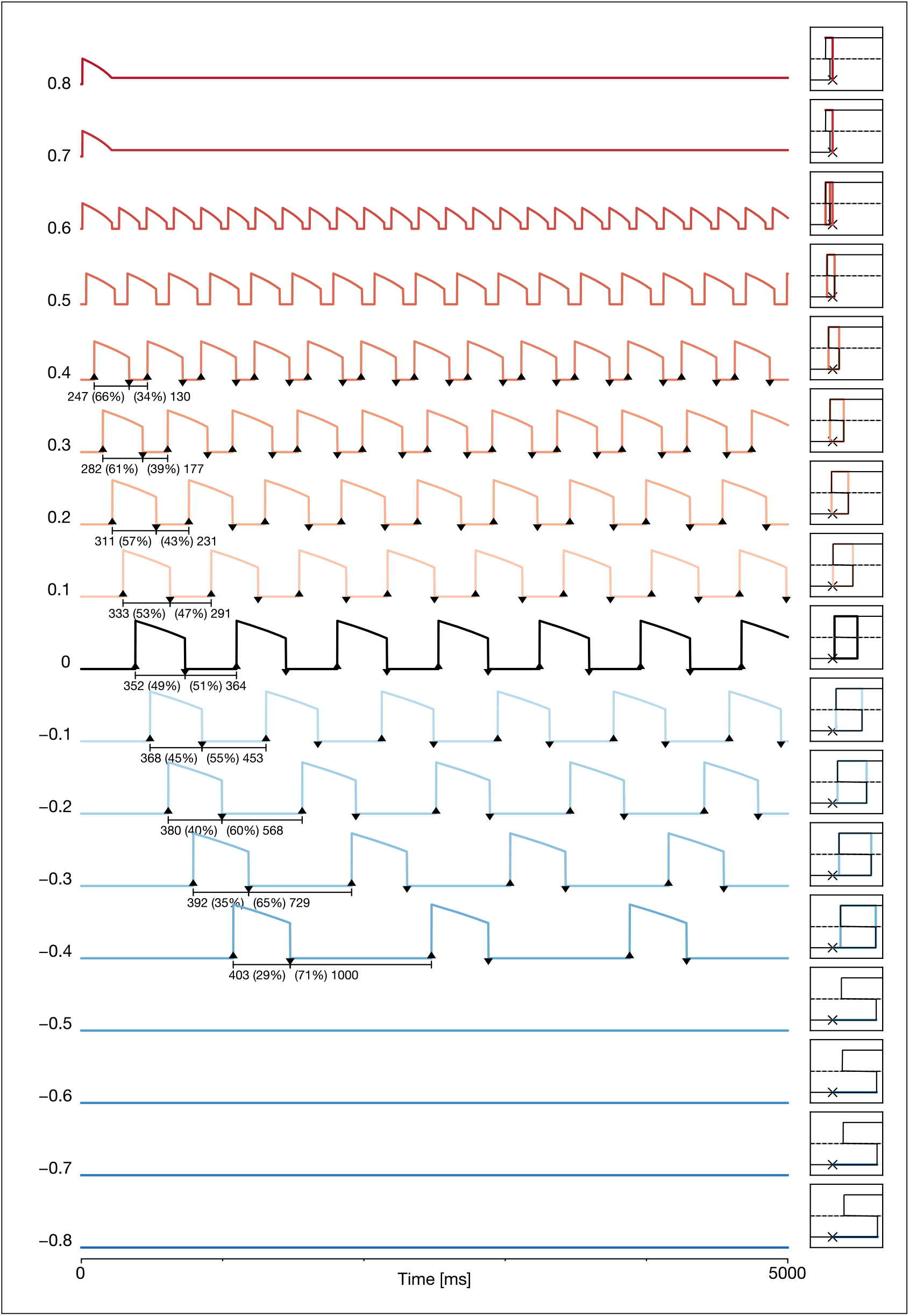
Simplified model response waveforms to constant stimuli. Activity for all stimulus strengths shown in ***Figure 3***, along with the corresponding phase portraits. Across stimulus strengths, the activity and phase portraits of the simplified model closely approximate that of the biophysical model. The simplified model decouples the shape of active ad quiet legs from the activity waveform’s shape and amplitude through the firing function.

**Figure 3—figure supplement 3.**
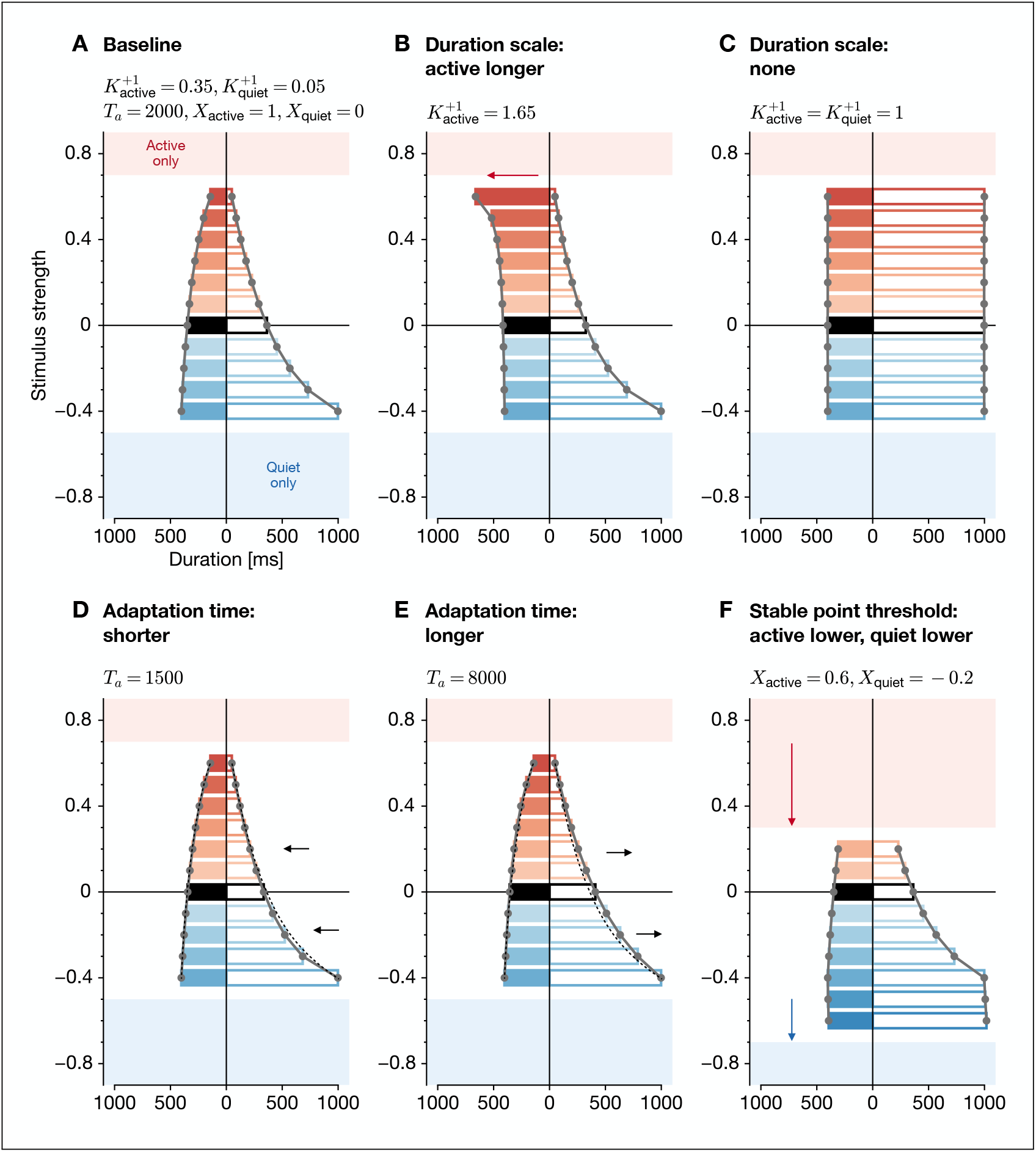
Simplified model effect of parameters on constant response. (**A**) Baseline response with default parameters from ***Table 2***, modified to use a rectangular burst shape (in order to remove possible confounding effects of the firing function on waveform measurements). (**B**) Increasing only the active duration scale makes the active duration 65% longer than the intrinsic active duration *T*_active_ at maximum input strength, while the quiet duration response is unchanged. (**C**) Removing both active and quiet duration scales results in the intrinsic durations *T*_active_ and *T*_quiet_ being expressed at all input strengths. (**D**) Shortening the adaptation time constant narrows the response profile. (**E**) Lengthening the adaptation time constant widens the response profile. (**F**) Lowering both active and quiet stable point thresholds lowers the input strengths that induce tonic and silent modes, respectively.

**Figure 4—figure supplement 1.**
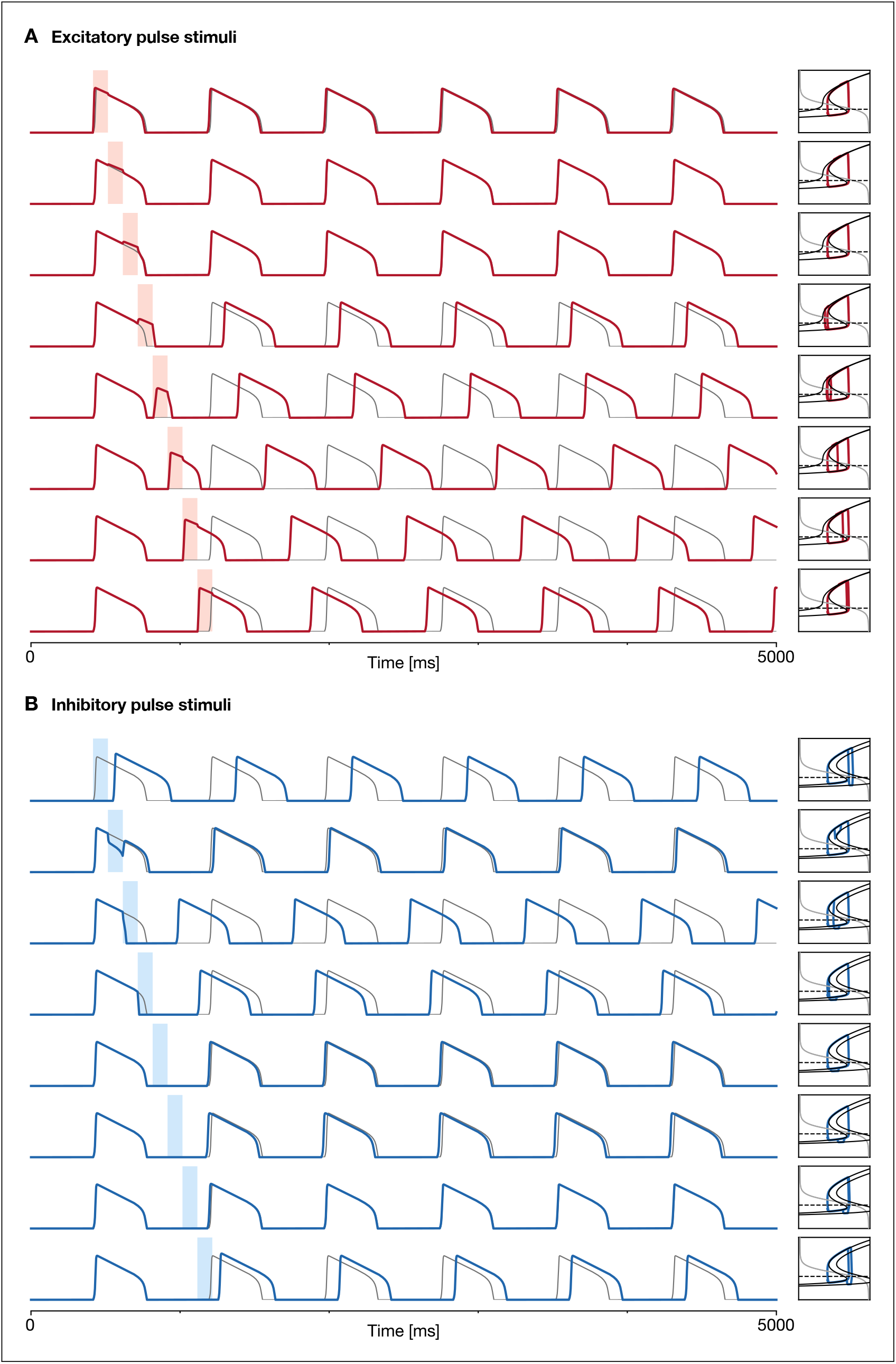
Biophysical model response waveforms to pulse stimuli. Activity for stimuli across various phases shown in ***Figure 4***, along with the corresponding phase portraits. (**A**) Excitatory pulses (strength 0.08, duration 100 ms) tend to lengthen the active phase and shorten the quiet phase. (**B**) Inhibitory pulses (strength –0.08, duration 100 ms) tend to shorten the active phase and lengthen the quiet phase.

**Figure 4—figure supplement 2.**
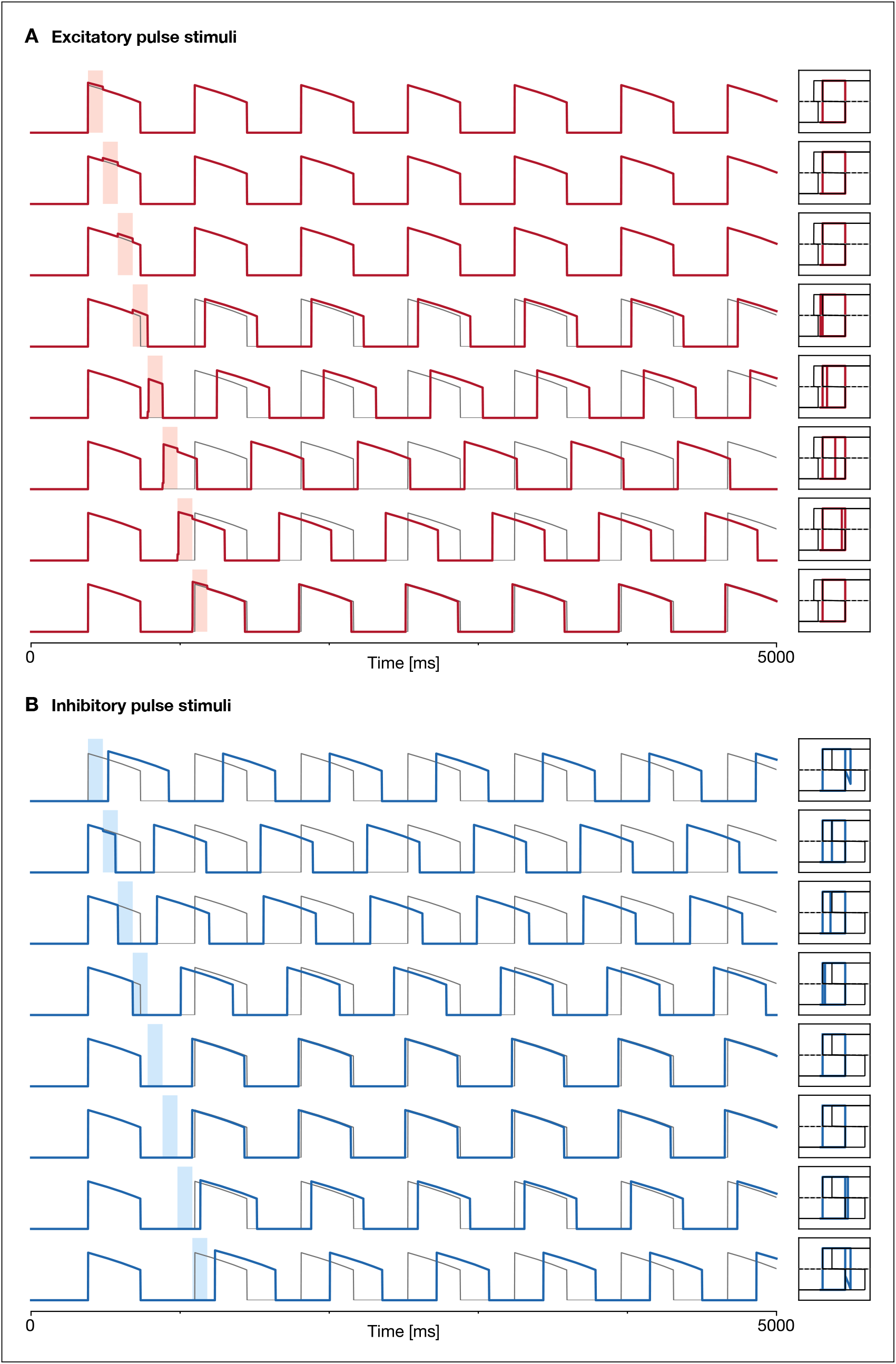
Simplified model response waveforms to pulse stimuli. Activity for stimuli across various phases shown in ***Figure 4***, along with the corresponding phase portraits. (**A**,**B**) Across both excitatory and inhibitory pulses, the activity and phase portraits of the simplified model closely approximate that of the biophysical model.

**Figure 4—figure supplement 3.**
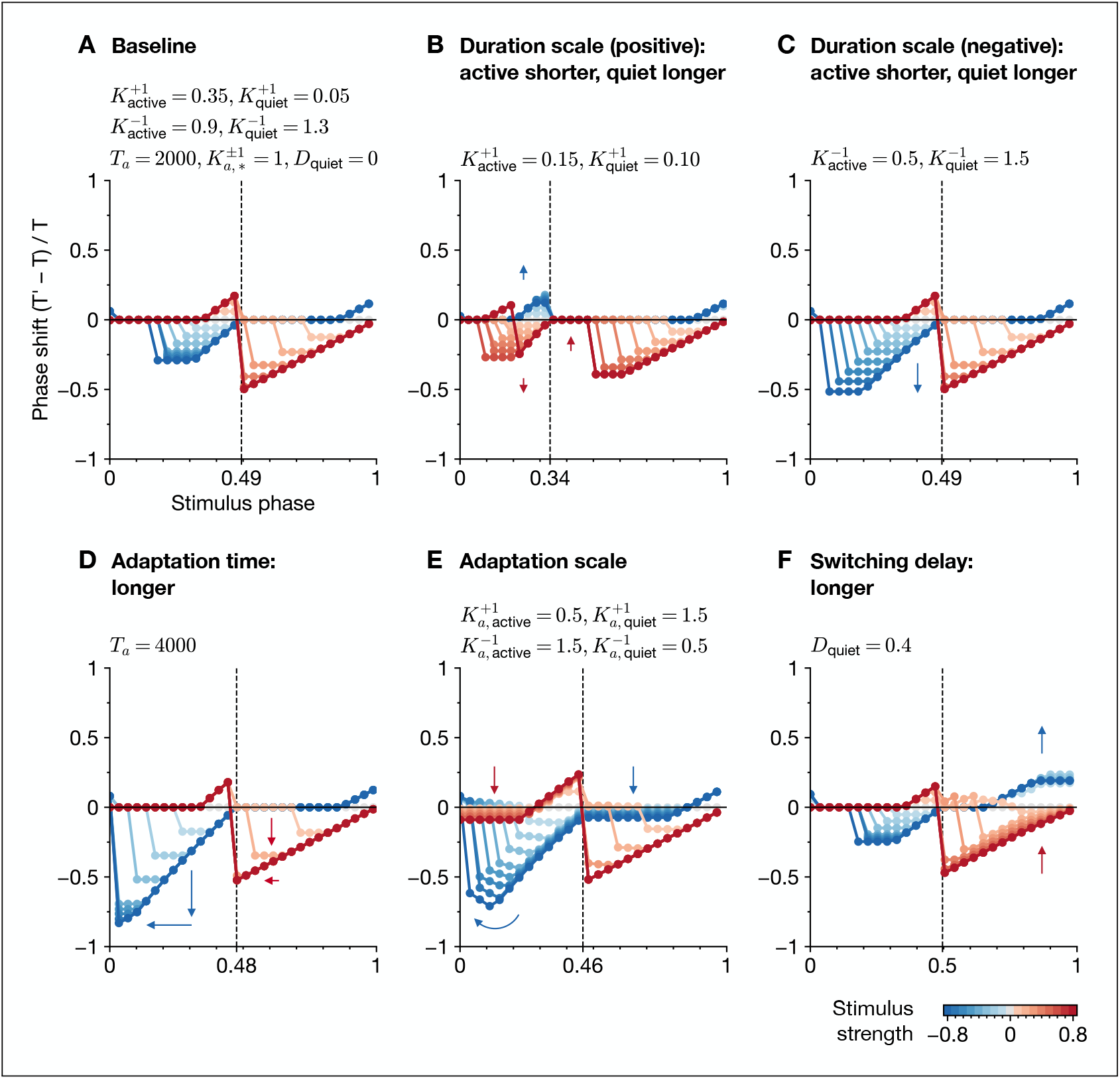
Simplified model effect of parameters on pulse response. (**A**) Baseline response with default parameters from ***Table 2***, modified to use a rectangular burst shape, no adaptation time constant scales, and no switching delay. (**B**) Changing the active and quiet duration scales (for positive input) results in different active and quiet bounds, which can dramatically alter the phase response curve in addition to the waveform duty cycle. In this example, excitatory and inhibitory pulses in the active phase have flipped advancing or delaying effects. (**C**) Shortening the active duration scale (for negative input) enables a stronger advancing effect by shortening the active phase. (**D**) Lengthening the adaptation time constant results in active and quiet bounds that are closer together (to maintain the desired active and quiet durations), with active and quiet nullcline legs overlapping less across different input strengths, leading to more responsive state switches due to falling off the legs. (**E**) Changing the adaptation time constant scales results in phase-independent speed changes along the active and quiet nullcline legs (corresponding to vertical translation of the flat segments in the phase response curve), while maintaining state switches due to falling off the legs (corresponding to the diagonal segments in the phase response curve). (**F**) Increasing the quiet switching delay creates a delay preceding the state switch, which is input-dependent since the delay is a constant fraction of the quiet duration.

**Figure 5—figure supplement 1.**
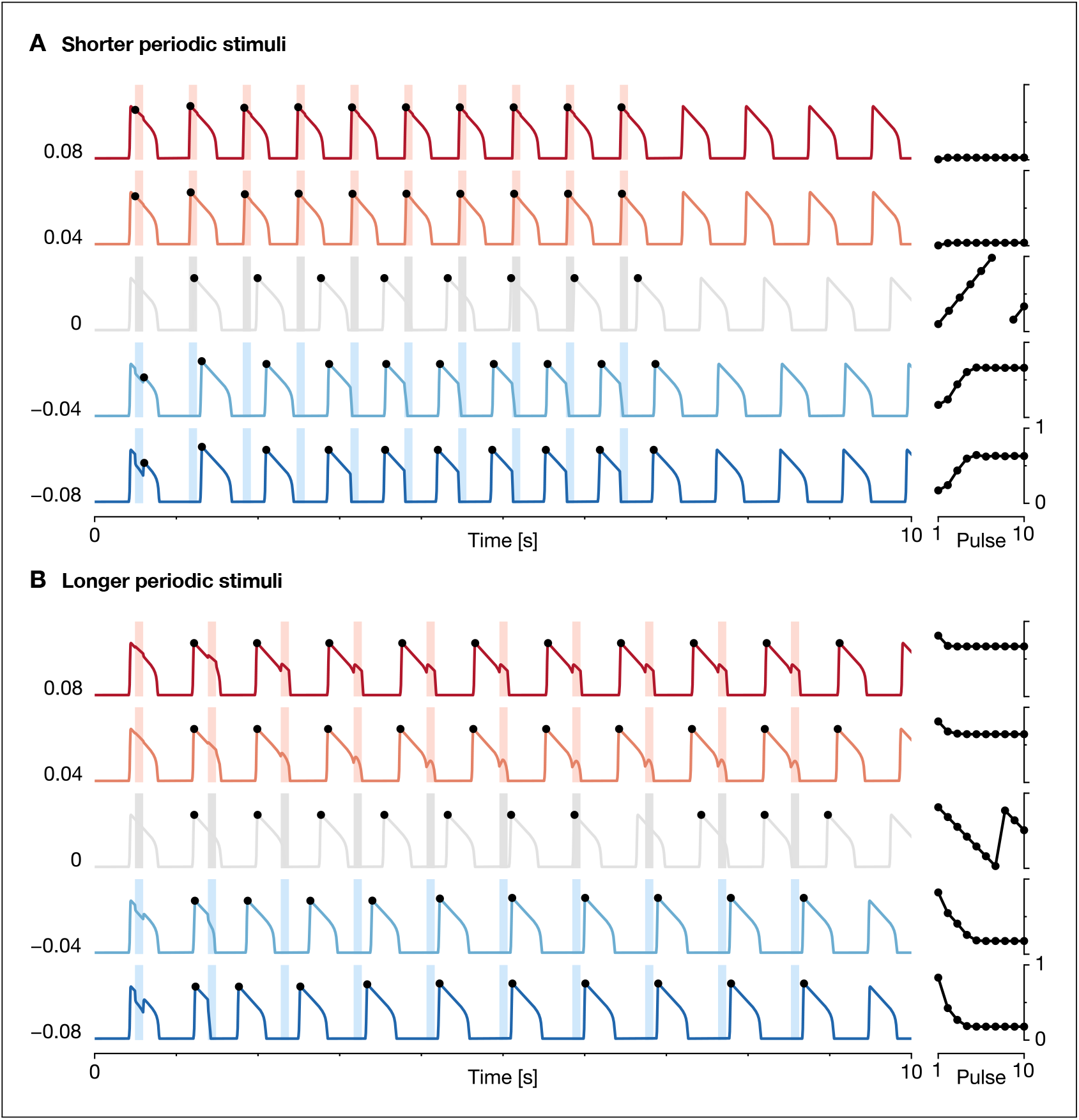
Biophysical model response waveforms to periodic stimuli. Activity for stimuli across various strengths and periods shown in ***Figure 5***, along with the corresponding phase plots. The entrainment response is closely related to the phase shift response from the pulse stimuli analysis. (**A**) Shorter-period pulse trains (normalized period 0.85, duration 100 ms) entrain leftward to stable points in the phase response curve (active start for excitatory pulses, quiet start for inhibitory pulses). (**B**) Longer-period pulse trains (normalized period 1.15, duration 100 ms) entrain rightward to stable points in the phase response curve (active end for excitatory pulses, quiet end for inhibitory pulses).

**Figure 5—figure supplement 2.**
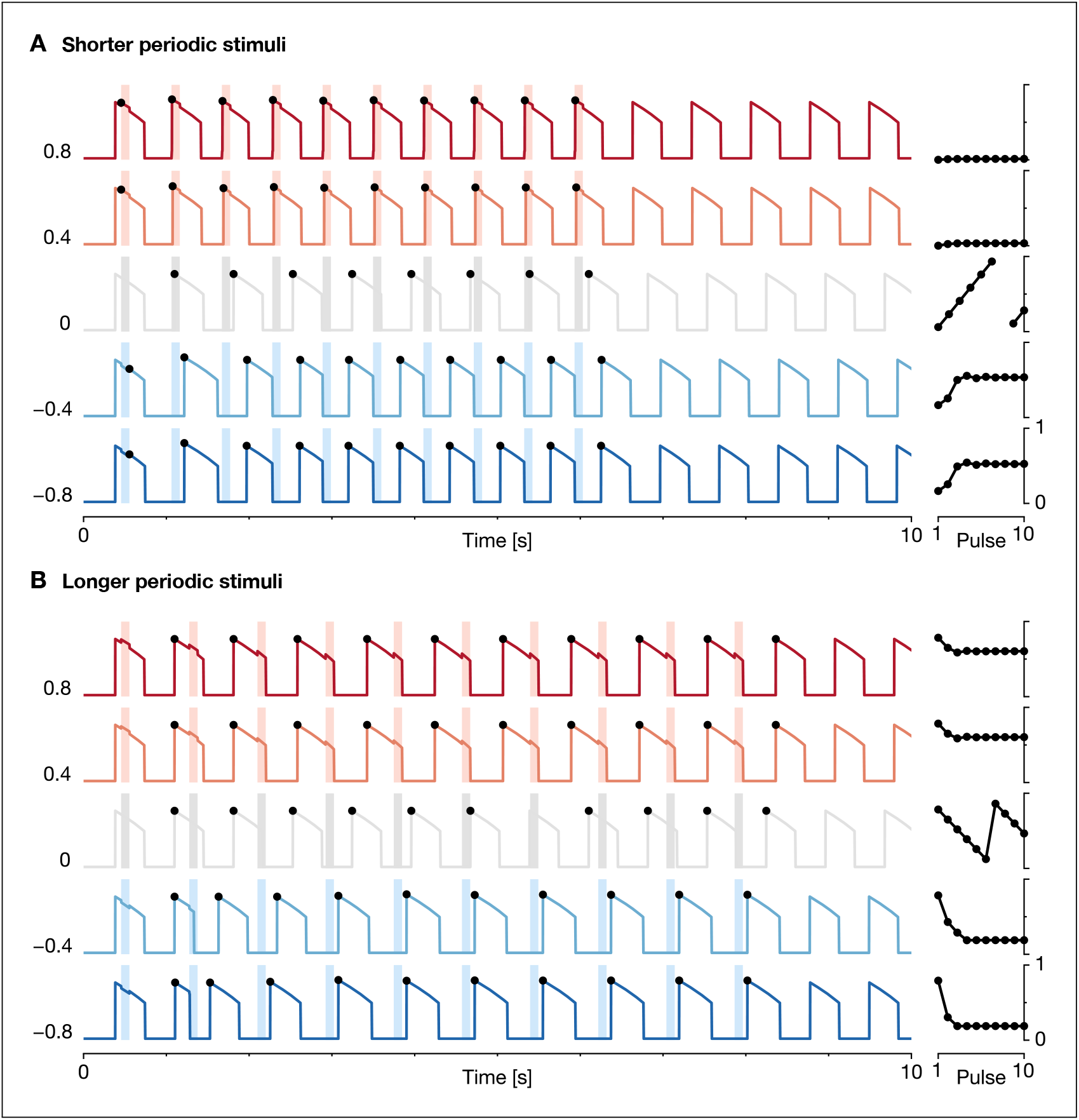
Simplified model response waveforms to periodic stimuli. Activity for stimuli across various strengths and periods shown in ***Figure 5***, along with the corresponding phase plots. (**A**,**B**) Across both shorter-period and longer-period pulse trains, the activity and phase plots of the simplified model closely approximate that of the biophysical model.

**Figure 5—figure supplement 3.**
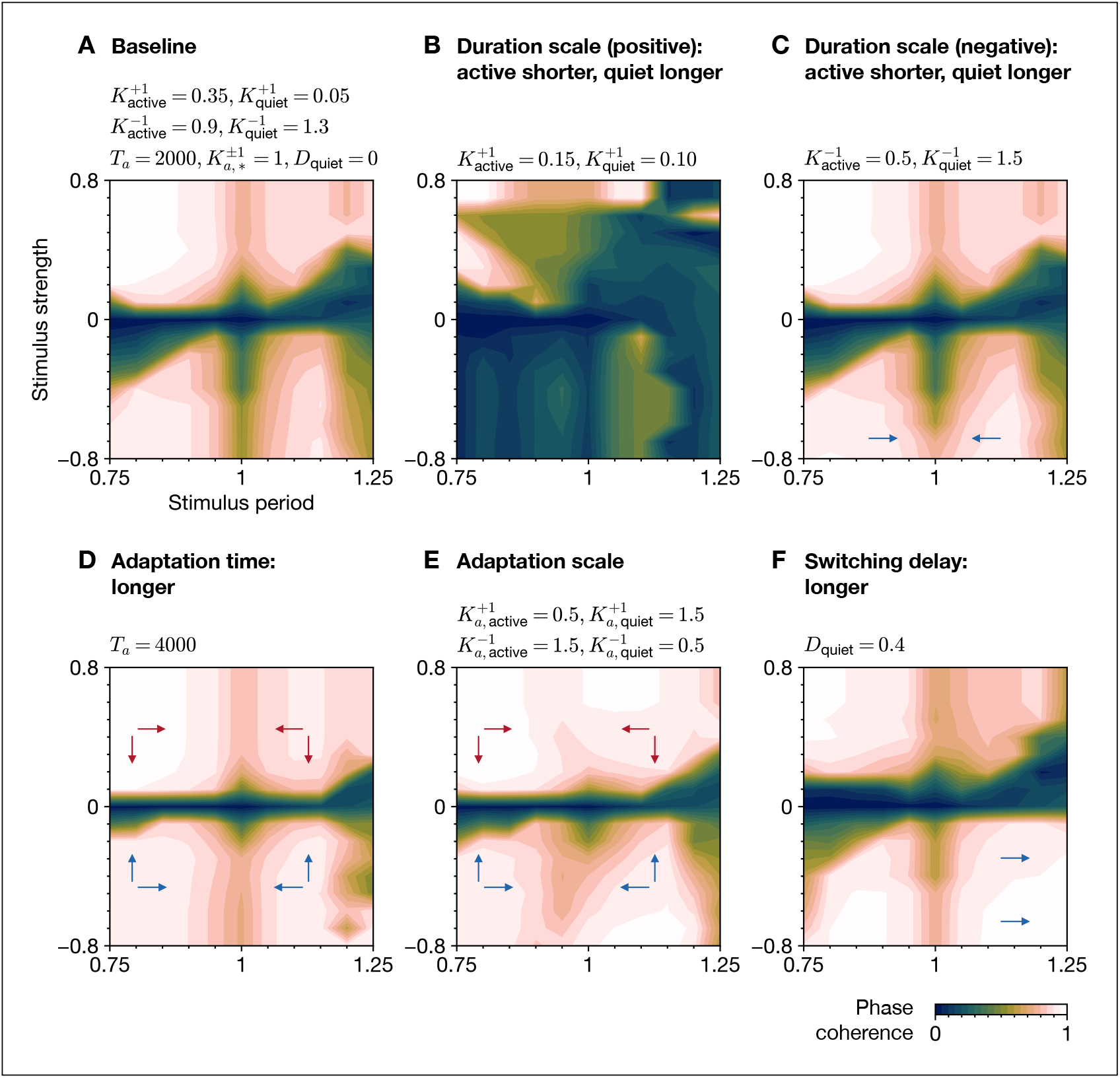
Simplified model effect of parameters on periodic response. (**A**) Baseline response with default parameters from ***Table 2***, modified as in the pulse stimuli analysis. (**B**) Changing the active and quiet duration scales (for positive input) results in different active and quiet bounds, which can dramatically alter the entrainment response. In this example, entrainment is difficult due to the lack of stable points in the phase response curve (positive-sloped zero crossings), except for a small region for positive input at the quiet-to-active switch that is utilized by shorter-period, excitatory stimuli. (**C**) Shortening the active duration scale (for negative input) increases entrainment for inhibitory stimuli due to the larger phase shift response for negative input. (**D**) Lengthening the adaptation time constant increases entrainment due to the larger phase shift response. (**E**) Changing the adaptation time constant scales increases entrainment due to the larger and more stable phase shift response. (**F**) Increasing the quiet switching delay increases entrainment for longer-period, inhibitory stimuli by shifting the stable point for negative input earlier and enabling a subsequent pulse to arrive in the stabilizing region of the phase response curve.

**Figure 7—figure supplement 1.**
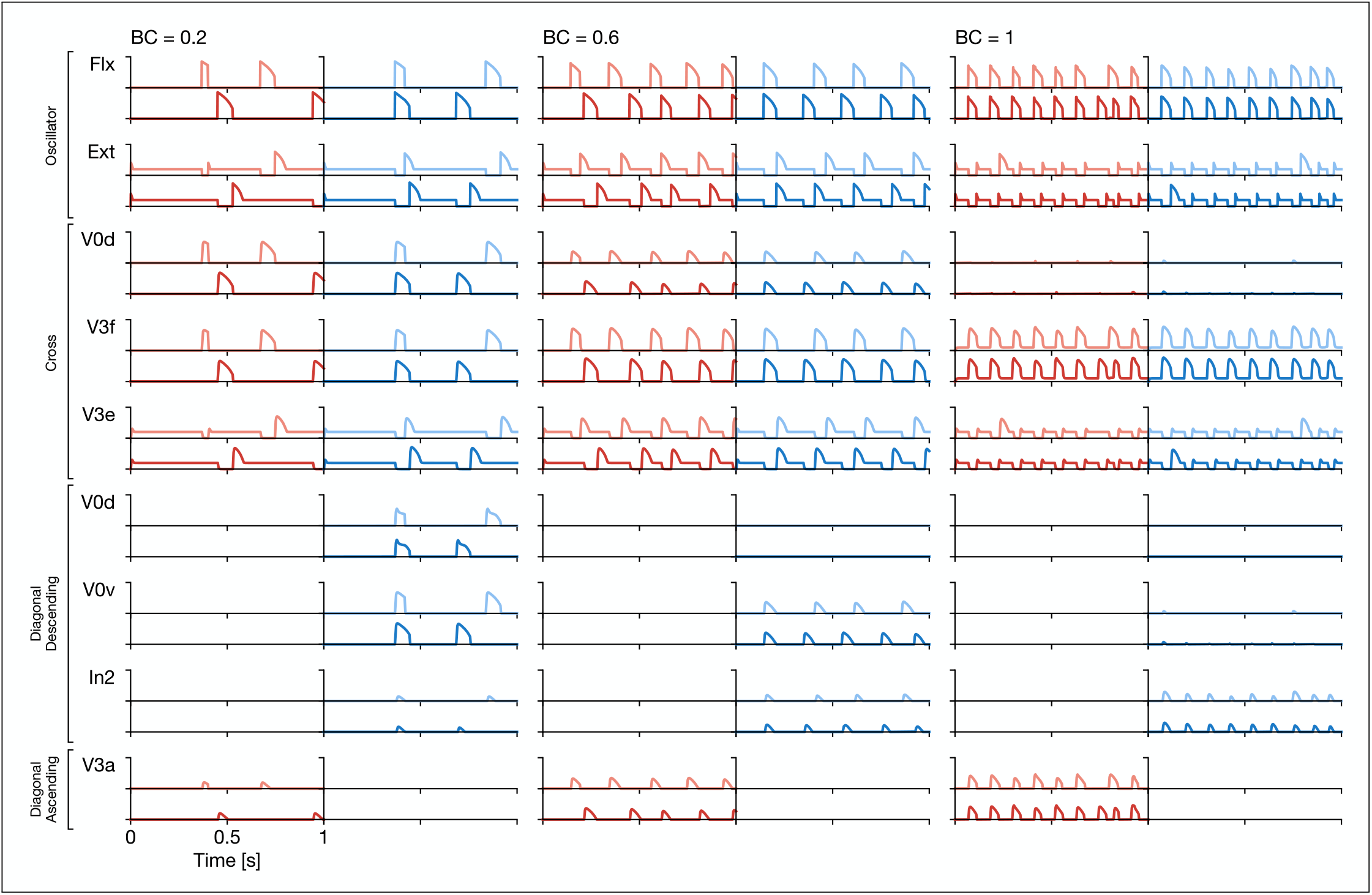
Activity of neuron types at different locomotor frequencies. Limbs are plotted with hindlimbs on the left (red) and forelimbs on the right (blue), with limb colors and neuron names corresponding to ***Figure 7***. Gaits are lateral-sequence walk (at BC = 0.2), trot (at BC = 0.6), and bound (at BC = 1). Flexor half-centers are nominally in silent mode and increase bursting frequency with BC. Extensor half-centers are nominally in tonic mode and increase burst frequency with BC. The V0d neurons promote limb alternation and dominate at walk (both cross and diagonal descending) and trot (cross only). The V3f neurons promote limb synchronization and dominate at bound (cross only). The V3e neurons promote limb synchronization and are important for establishing the correct phase relations for lateral-sequence walk by suppressing rebound (***Danner et al., 2016***). The V0v neurons and V3a promote limb synchronization and dominate at trot (diagonal descending and diagonal ascending, respectively). These activity profiles approximate those from ***Zhang et al. (2022)***, which modeled the circuit using populations of conductance-based spiking neurons.

## Notes

### Competing Interest Statement

The authors have declared no competing interest.

